# A latent 5-dimensional space for action representation: Geometric validity and dissociation from kinematics

**DOI:** 10.64898/2026.06.09.731166

**Authors:** Nick E. Barraclough

## Abstract

A fundamental question in visual perception is how the human visual system transforms the rich kinematic information present in observed actions into coherent social meaning. We addressed this by validating a 5-dimensional action space model - defined by Formidableness, Friendliness, Locomotion, Abduction, and Environmental Interaction - as a perceptual representation of avatar-conveyed actions. Using Representational Similarity Analysis, we first demonstrated strong topological correspondence between the model’s geometry and the structure of independent perceptual judgements, with cross-validated regression confirming the model as a generative framework that reliably predicts how observers evaluate novel actions. A morphing paradigm further revealed that perceptual ratings scaled approximately linearly with geometric distances along model dimensions, with each dimension selectively predicting its corresponding perceptual quality, satisfying the criteria for a valid psychological metric space. Critically, the 5D model showed substantially stronger alignment with semantic representations of actions than with their raw skeletal kinematics - an association robust to statistical control for kinematic similarity. This dissociation suggests that higher-order social-evaluative dimensions of action perception are largely invariant to low-level motion statistics, consistent with a hierarchical visual processing architecture in which kinematic input is progressively abstracted into a compact, semantically organised representational space optimised for social inference.

## Introduction

Action specific information derived from observing body posture and movement constitute critical social cues that influence behaviour regarding interpersonal interactions (Ansuini et al., 2015; Becchio et al., 2012; Becchio et al., 2010; Sartori et al., 2011). For example, observers readily infer force (Alaerts et al., 2010; Runeson & Frykholm, 1981) emotions (de Gelder, 2006; de Gelder et al., 2010), gender (Troje et al., 2006), dominance (Carney et al., 2005), trustworthiness (Koppensteiner et al., 2016), intentions (Cole et al., 2017; de Lange et al., 2008), identity (Troje et al., 2005) and personality traits (Koppensteiner, 2013; Thoresen et al., 2012) of other people from action information.

Despite the social utility of these judgements, only recently have attempts been made to identify the organising principles underpinning how this information is represented. Emerging evidence suggests that actions are represented within a multidimensional ‘action space’ where different actions are represented as coordinates within a continuous geometric structure (Barraclough & Wightman, 2026; Bockes et al., 2025; Dima et al., 2022; Kabulska & Lingnau, 2023; Tucciarelli et al., 2019; Vinton et al., 2023). These models draw on the concept of a ‘conceptual space’ (Gärdenfors, 2004a, 2004b). This framework proposes that for different stimulus domains (e.g. colour, odours, face traits, actions etc.) that the distance between items within the representational space reflects their perceived similarity (Shepard, 1962a, 1962b). Within a representational space, the principal dimensions define the meaningful qualities on which we make judgments about items within the domain. Consequently, the coordinates of an item within the space indicate how well it conveys the different qualities defined by the space dimensions. For example, for an action located within the 4-dimensional action space model described by Vinton et al. (2023), an action’s co-ordinates can describe its friendliness, formidableness, intentionality, and body abduction (Vlasceanu et al., 2024).

For many stimulus domains, it has been shown that the structure of the representational space determines the judgments we make about items in the domain. This has included domains for shape (Nosofsky, 1986; Shepard et al., 1961), colour, (Shepard, 1962a), objects (Edelman, 1998), musical pitch (Shepard, 1982), faces (Nosofsky, 1991; Valentine, 1991) and word meaning (Landauer & Dumais, 1997), demonstrating the psychological validity of these dimensional models. This relationship has been tested in a number of ways.

First, dimensional models should be both robust under methodological convergence and reflect the very qualities and similarities that humans perceive in the real world (Indow & Kanazawa, 1960; Wiggins, 1979). For example, recently Thornton and Tamir (2020) validated a 3-dimensional model of mental state representation (defined by rationality, social impact, and valence) by demonstrating a structural match between their model and human judgments. Using Representational Similarity Analysis (RSA; Kriegeskorte & Kievit, 2013; Kriegeskorte et al., 2008), they showed that the distances between states in the 3D space significantly predicted the similarity patterns found in both neural activity and behavioural ratings. Furthermore, using cross-validated regression they showed that the model serves as a generative framework, where each item’s coordinates (location) within the space could reliably predict how participants would attribute qualitative features to those states in held-out data. Similar predictive approaches have been successfully applied in other visual domains, notably for faces. Using RSA, Stolier et al. (2018) showed that the conceptual structure of trait space determines the perceptual representation of faces. Likewise, Jozwik et al. (2022) used RSA and cross-validated predictive modelling to confirm that geometric distances within a statistically defined face space capture the similarity relationships humans perceive among individual faces. This is consistent with computational models of visual systematicity (e.g. Edelman & Intrator, 2003), which suggest that human perception relies on graded, continuous representational spaces rather than discrete categories. Edelman and Intrator (2003) thus provide a formal computational basis for the idea that an item’s specific coordinates within a representational space determine the qualitative attributes humans perceive in it. Together, these studies suggest that representational spaces can function as universal, generative frameworks that govern human judgments across domains both physical and social.

Second, in true perceptual representational spaces, changes in subjective qualities scale systematically, and approximately linearly, with distances in that space; metric distances correspond to perceptual differences. In an early example of this principle Indow and Kanazawa (1960) showed that Euclidean distances in a derived colour space mapped linearly onto perceived differences in hue and saturation. Subsequently, this metric validity has been confirmed in complex biological and social domains. For example, Troje (2002) used a morphable model of human gait to show that human motion can be decomposed into a linear ‘gait space,’ where the geometric distance along specific vectors reliably and selectively predicts the intensity of perceived qualities such as gender or weight. Similarly, in the face domain, Jozwik et al. (2022) used a morphable face model to demonstrate that representational distances within a statistically defined space capture the dissimilarity relationships humans perceive among individuals. These findings align with Shepard’s (1987) *Universal Law of Generalization*, which suggests that psychological distance governs the relationship between stimuli. Collectively, these studies imply that for a dimensional model to be a valid representation of the domain, metric distances between two points in that space must selectively and linearly predict the qualitative differences humans perceive in the underlying items.

Drawing on this previous research across a multitude of different perceptual domains, in this paper we validate a recent multidimensional model of action space (Barraclough & Wightman, 2026). Using an EFA-to-CFA approach, Barraclough and Wightman (2026) showed that evaluations of the characteristics of human actions are best modelled by a 5-dimensional action space defined by the fundamental qualities of action Formidableness, Friendliness, Locomotion, Abduction, and Environmental interaction. This model is a recent development of the Vinton et al.’s (2023) 4D action space model defined by the very similar dimensions of Formidableness, Friendliness, Intentionality, and Abduction. The updated 5D model incorporates additional data to resolve differences between earlier competing action space models (Bockes et al., 2025; Dima et al., 2022; Kabulska & Lingnau, 2023; Tucciarelli et al., 2019; Vinton et al., 2023). Consequently, we tested whether the dimensions determined by Barraclough & Wightman’s (2026) latent factor modelling are ‘psychologically real’ - in that they correspond with psychologically meaningful perceptual qualities that guide the way we make judgments about the actions of other individuals.

A fundamental challenge in understanding action representation lies in distinguishing between the perceptual representation of physical action form and motion (Giese & Poggio, 2003) and the cognitive representations of the social meaning of actions (Van Overwalle & Baetens, 2009). While the human visual system is tuned to the kinematics of biological motion (Johansson, 1973), the ultimate goal of observation is to infer the underlying social affordances of the actor (Becchio et al., 2010; Friston et al., 2011; Moore & Haggard, 2008; Sartori et al., 2011). If Barraclough & Wightman’s (2026) 5D action space represents a higher-order cognitive map, it should exhibit a degree of kinematic invariance. This requirement is particularly critical given the rise of Spatio-Temporal Graph Convolutional Networks (ST-GCNs), which achieve high action classification accuracy by optimizing for raw kinematic patterns (Yan et al., 2018) but frequently fail to capture the nuances of human social perception (Garcia et al., 2025; Garcia et al., 2024). This ‘alignment gap’ suggests that a valid model of action space must capture the specific social-evaluative contribution that remains elusive to purely kinematic computational models.

In this study we first assessed the psychological validity of the 5D action space model in two experiments. In Experiment 1 we tested the perceptual validity of the hypothesized 5D action space. We used RSA to compare a factor-based Representational Dissimilarity Matrix (RDM) against a behavioural RDM derived from participant ratings, allowing us to assess both the convergent and discriminant validity of the five dimensions. In line with the principle that psychological geometry governs subjective judgment (Shepard, 1987), we further tested the model’s predictive power using a cross-validated regression analysis. We predicted that an action’s specific coordinates (or location) within this 5D space would reliably determine the degree to which participants attribute different qualities to that action.

In Experiment 2 we tested whether the dimensions of the 5D action space behave as perceptual metrics. We generated morph continua between exemplar actions, creating novel intermediate actions whose model-derived coordinates varied smoothly between the two endpoints. By drawing upon Shepard’s (1987) Universal Law of Generalization, we aimed to test the logic that systematic changes in geometric distance within the model should translate into proportional changes in perceived qualities – characteristic of a valid psychological space. Participants then rated these morphed actions on each of the five action qualities to determine if the predicted factor scores were reflected in monotonic and approximately linear selective changes in the corresponding perceptual ratings. Consistent linear scaling would demonstrate that distances in action space correspond to proportional differences in perception, thereby validating the model’s metric structure.

Finally, we formally tested the degree to which the dimensions of the 5D action space are related to action structural kinematics and semantic information in Experiment 3. We used RSA to compare between RDMs of the 5D action space and its individual dimensions, and RDMs of the kinematics and semantic information of the actions. If dimensions represent higher-order cognitive properties, they should exhibit a degree of kinematic invariance. That is, while a representation must be grounded in physical reality to be useful, it should dissociate from the raw skeletal statistics to prioritize more conceptual social-evaluative information. By comparing the relative similarity of action space dimensions to both lower- and high-level references, we quantify the hierarchical position of each dimension within the social visual processing network (cf. McMahon et al., 2023).

Although the aim of this study is to evaluate the 5D model of action space we additionally provide in the supplemental materials identical computations for the 4D action space model (Vinton et al., 2023) from which it developed. A comparison of analyses of the earlier and later models confirmed equivalent psychological and metric validity and dissociation from raw skeletal kinematics.

## Experiment 1 – perceptual validity

### Methods

#### Participants

Two hundred and twelve participants completed the rating experiments. Sixty-one of these were removed from the data analysis as they failed to respond correctly to 75% or more of the catch trials (see procedure). This ensured we had a minimum of thirty participants rating all actions on each of the action qualities. A power analysis was not appropriate in this instance as we did not conduct inferential statistics on the data; however, the central limit theorem states that a sample size of 30 would be appropriate for most distributions. Demographic information of those who completed each rating task are described in Table 1. Participants were recruited via the internal University of York SONA system and compensated with course credit. All participants reported normal or corrected-to-normal vision. This experiment was approved by the ethics committee of the Department of Psychology, University of York, and was performed in accordance with the ethical standards laid down in the 1964 Declaration of Helsinki.

**Table 1.** Participant demographic information.

*Participant demographic information*
| Characteristic | N | Gender | Mean age | SD age |
| --- | --- | --- | --- | --- |
| Formidableness | 30 | F=26, M=4 | 19.03 | .81 |
| Friendliness | 30 | F=24, M=5, Other=1 | 19.00 | .59 |
| Locomotion | 30 | F=27, M=2, Other=1 | 20.03 | 2.48 |
| Abduction | 31 | F=25, M=6 | 19.35 | 1.70 |
| Environmental Interaction | 30 | F=25, M=4, Other=1 | 19.57 | 1.77 |

#### Stimuli

Stimuli were the actions from Barraclough and Wightman (2026) and Vinton et al. (2023) and are available from the Open Science Framework at https://osf.io/4vew8/. Actions were conveyed by an androgynous volumetric avatar (see Figure 1) to isolate action information from body posture and body movements from other sources of information about the person (e.g. face, body morphology, clothing etc.) that are naturally confounded in photorealistic videos of human actions. In this prior study (Barraclough & Wightman, 2026) the loading of 240 different actions onto 5 different latent dimensions (Formidableness, Friendliness, Intentionality/Planned, Abduction) was calculated using Exploratory Factor Analysis (EFA). In this framework, action factor scores are equivalent to coordinates within the 5-dimensional geometry.

**Figure 1.**
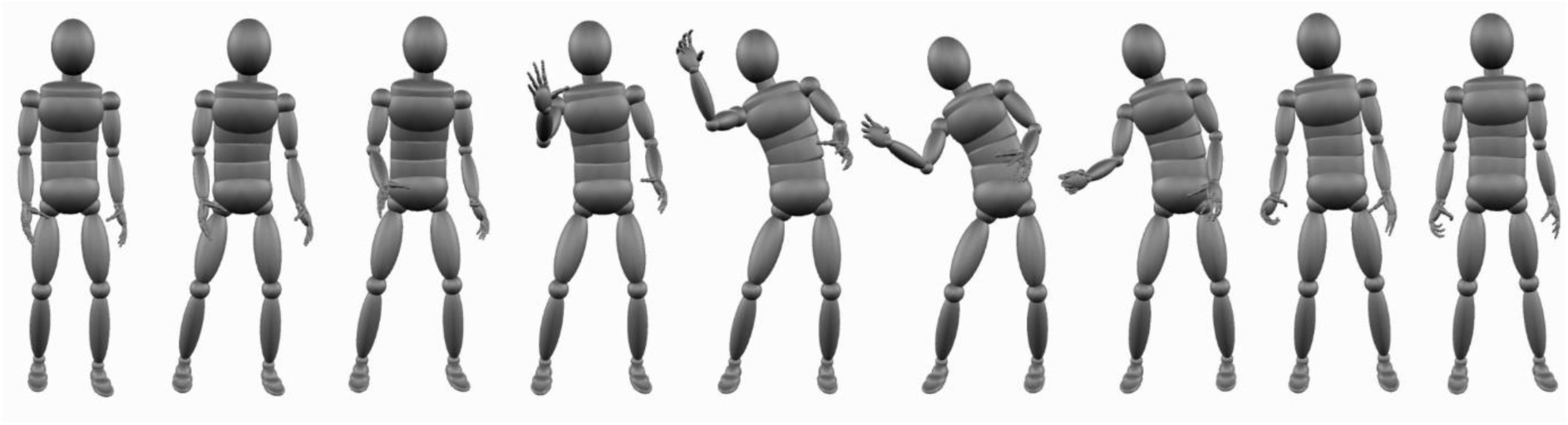
Illustration of frames (1, 13, 25, 37,49, 61, 73, 85, 97) from the catching action (from Barraclough & Wightman, 2026; Vinton et al., 2023).

#### Procedure

All rating experiments were implemented within the Gorilla platform (Anwyl-Irvine et al., 2021; Anwyl-Irvine et al., 2020). Once participants entered the site via an internet browser (Chrome, Firefox or Safari) on a laptop or desktop PC, participants completed a consent form, recorded their age and gender, and then instructions on the experimental task were displayed. Initially participants took part in a practice block of 8 trials to familiarise them with the nature of the action stimuli, the task they were to perform, and the pacing of the experiment. On each trial they first observed a fixation cross in the centre of a white screen for 500 ms, then the video of the action for its duration, and finally a response screen showing a 1-9 Likert scale where participants had to indicate their rating of the action (e.g. Friendliness) by clicking an onscreen button with the mouse. Once a response was registered, the next trial commenced. If participants failed to respond within 2 s of the end of the action, a prompt “Please respond faster!” was shown on screen to encourage participants to indicate their immediate first impressions of the action. Following completion of the practice block, participants started the main experiment. The experiment consisted of 244 trials in total, in 240 trials one of the 240 different actions were shown, in 4 catch trials the response screen explicitly asked the participant to press a specific button, for example “Press 3”. These catch trials were used to assess whether participants were paying attention during the experiment; participants were eliminated from analysis if they failed to respond correctly to 75% or more of the catch trials. The experiment was divided into 4 blocks of 61 trials, and participants were given the opportunity to take breaks of up to 2 minutes to help maintain concentration through the duration of the experiment. A progress bar was presented on screen along with the response buttons to provide participants with an indication of how far through the experiment they were; on average each experiment took approximately 30 minutes to complete. Once the experiment completed, participants were shown onscreen debriefing information.

For those rating actions on formidableness, the instructions were to: “rate the actions on how formidable they appear on a scale from 1 to 9. With: 1 = Very feeble, 5 = Neither feeble nor formidable, 9 = Very formidable. For example, punching someone hard might be seen as very formidable, moving an object slightly might be seen as very feeble, whilst opening a door might be seen as neither feeble nor formidable.” For those rating actions on friendliness, the instructions were to: “Rate the actions on how friendly they appear on a scale from 1 to 9. With 1 = Very unfriendly, 5 = Neither unfriendly nor friendly, 9 = Very friendly. For example, waving at someone might be seen as very friendly, punching someone might be seen as very unfriendly, whilst opening a bottle might be seen as neither unfriendly nor friendly.” For those rating actions on locomotion, the instructions were to: “Rate the actions on how much the action involves the coordinated movement of the whole body to change the actor’s physical location on a scale from 1 to 9. With: 1 = Prevents the change of actor location, 5 = Neither prevents nor changes the actor’s location, 9 = Very much changes the actor’s location”. For those rating actions on abduction, the instructions were to: “Rate the actions on how abducting they appear on a scale from 1 to 9. With: 1 = Very adducting (towards body), 5 = Neither adducting nor abducting, 9 = Very abducting. For example, kicking a ball might be seen as very abducting (movement away from the body), picking an object up from the floor might be seen as very adducting (movement towards the body), whilst making a phone call might be seen as neither adducting nor abducting. For those rating actions on environmental interaction, the instructions were to: “Rate the actions on how much the action is directed towards an inanimate object and changing the physical environment on a scale from 1 to 9. With: 1 = Definitely not towards the physical environment, 9 = Definitely towards the physical environment. For example, dancing might be seen as not changing the physical environment, whilst cutting a loaf of bread with a knife might be seen as changing the physical environment.

### Results

For each of the 5 action qualities, interclass correlation coefficients (ICC; Koo & Li, 2016) were calculated to assess the consistency of ratings between participants. ICC estimates and their 95% confident intervals were calculated using SPSS statistical package version 29 (SPSS Inc, Chicago, IL) based on mean-rating (k = 30 or k = 31), consistency, 2-way random-effects models. In addition, Cronbach’s alphas were calculated to assess the internal consistency of each rating scale. These results are presented in Table 2. All action quality ratings were found to have moderate or greater reliability, and so all were included in subsequent analyses.

**Table 2.**
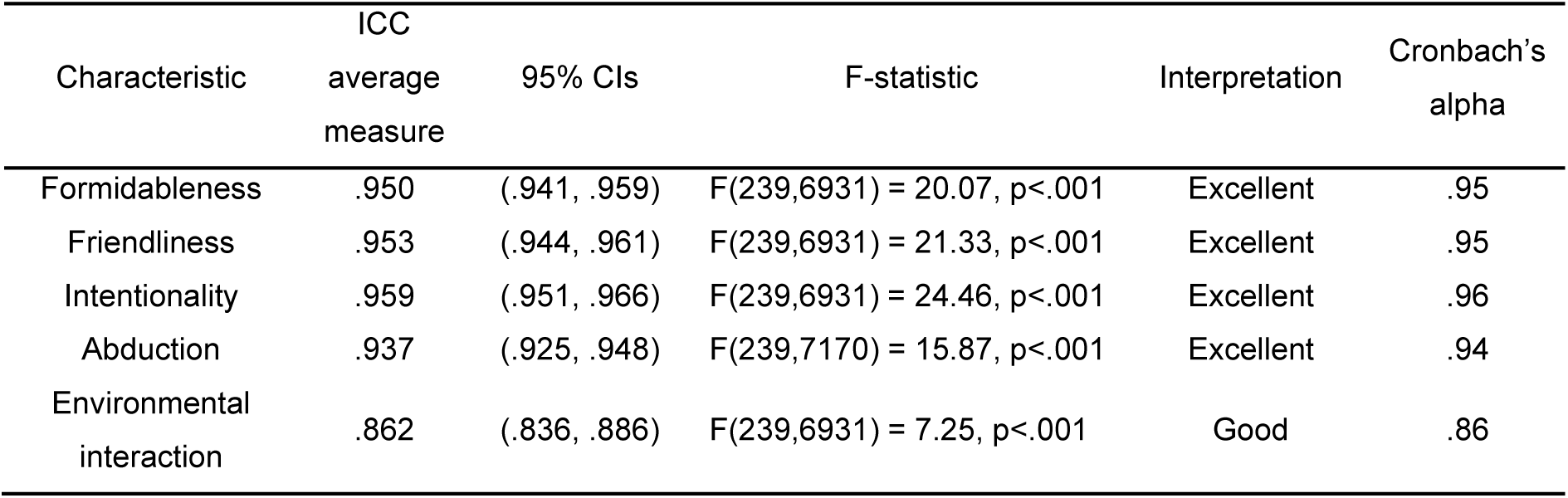
Interclass correlation coefficients (ICC) and Cronbach’s alpha for all 5 action qualities.

To examine how the structure of the average ratings of the different actions (rating space) relates to the structure of factor scores (factor space), we first calculated separate representational dissimilarity matrices (RDMs; cf. Kriegeskorte et al., 2008) for the factor scores and action ratings (left and middle panels respectively in Figure 2). To visualise their relationship, we plotted pairwise distances between all actions in rating space against the corresponding distances in factor space (Figure 2 right panel). There is a clear positive relationship (Spearman’s r = .81, p = .0001), with more distant action pairs in factor space generally corresponding to more distant pairs in rating space This correspondence was formally assessed using a Mantel test. The rows and columns of the rating RDM were randomly permuted 10,000 times, and the Spearman correlation with the factor RDM was computed for each permutation to generate a null distribution of correlations. The p-value was computed as the proportion of permuted correlations exceeding the observed correlation. The Mantel test confirmed a strong relationship between rating-derived and factor-derived RDMs (Spearman’s r = .79, p=.0001) indicating that the geometry of the 5D factor space predicted judgments of action qualities.

**Figure 2.**
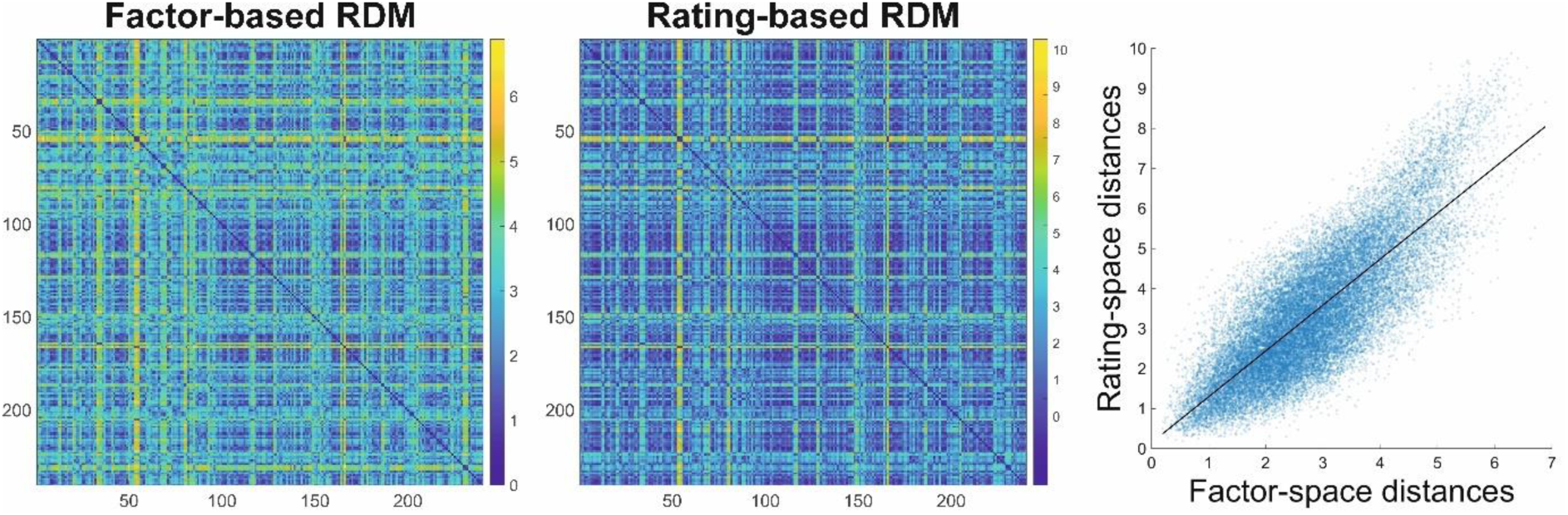
Representational dissimilarity matrices Note: The left panel illustrates the dissimilarity matrix for actions derived from CFA-EFA factor scores (Barraclough & Wightman, 2026). The middle panel illustrates the dissimilarity matrix for actions derived from ratings of actions against different qualities. The right panel illustrates the relationship between distances between pairs of actions (each marker) in rating space and factor score space; black line indicates the best-fit regression line.

Next, we characterised the interrelationships among the 5 factors. The original model was derived using an oblique rotation, which allowed factors to correlate, and consequently the latent factor correlation matrix showed moderate inter-factor associations (Table 5 in Barraclough & Wightman, 2026). To better visualise how the 240 actions were distributed in the derived action space, we also computed correlations between factor scores (Figure 3, left panel). As expected, given the oblique rotation, inter-factor correlations were moderate in size (rs .04 to .56).

**Figure 3.**
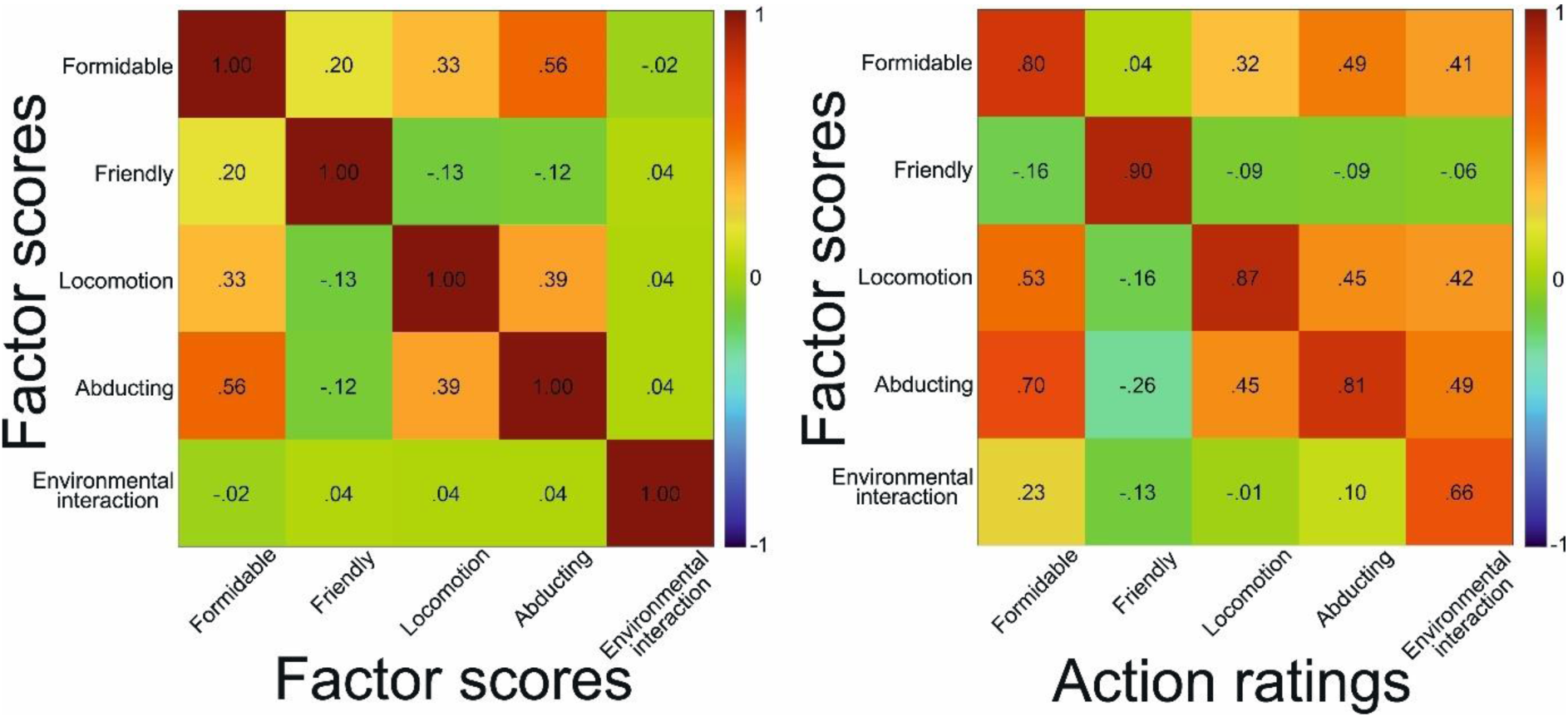
Action quality correlation matrices Note: Left panel: factor score correlation matrix. Right panel: action ratings correlated with factor scores.

To assess convergent validity, we correlated factor scores with ratings of corresponding action qualities; these were strong (rs .66 to .90; Figure 3, right panel), indicating each factor maps closely onto the subjective evaluations of its intended quality. Cross-dimension correlations were generally weaker, although with some large secondary associations (e.g., ratings of abduction correlated with formidableness, r = .70). For discriminant validity, multiple regression and variance partitioning analyses showed that in each case the matched factor was the strongest predictor of its rating (βs = .65 to .91). Unique R^2^ values (.27 to .72) confirmed that each factor explained variance above and beyond shared contributions from other, correlated factors (Table 3; see Table S1 in the supplemental materials for full regression results). Together, these results demonstrate both convergent and discriminant validity of the 5D action space dimensions, supporting their interpretation as distinct but related qualities of actions.

**Table 3.** Regression and variance partitioning for ratings predicted by factors.

| Rating (DV) | Target factor (IV) | $\beta$<br>(target) | Full model<br>$R^2$ | Unique $R^2$<br>(target) | Largest cross-predictor<br>$\beta$ | Shared<br>$R^2$ | VIF |
| --- | --- | --- | --- | --- | --- | --- | --- |
| Formidableness | Formidableness | .69 | .88 | .27 | Environmental interaction<br>$\beta = .22$ | .44 | 1.69 |
| Friendliness | Friendliness | .91 | .86 | .72 | Formidableness: $\beta = -.25$ | .09 | 1.16 |
| Locomotion | Locomotion | .84 | .78 | .56 | Environmental interaction<br>$\beta = .22$ | .19 | 1.24 |
| Abduction | Abduction | .74 | .69 | .33 | Environmental interaction<br>$\beta = .25$ | .33 | 1.66 |
| Environmental interaction | Environmental interaction | .65 | .73 | .41 | Formidableness: $\beta = -.24$ | .21 | 1.01 |

Finally, using a cross-validated regression analysis, we tested whether the 5D model of actions can be used to predict how participants rate action qualities. Each rating dimension was predicted from the factor scores for the 5 dimensions using a 10-fold cross validation procedure. Here the dataset of 240 actions was split into 10 partitions, models were then trained on 90% of the data and tested on the remaining 10%. This was repeated across folds. The correlation between predicted and observed ratings and the corresponding cross-validated R^2^ values were used to determine predictive accuracy. To assess whether the obtained predictive correlations exceeded chance, a permutation test with 1000 iterations was conducted in which the correspondence between factor scores and ratings was randomly shuffled. P-values were calculated as the proportion of permutations yielding a correlation >= than the observed one.

Predictive performance was high across all five rating dimensions (see Table 4). Cross-validated predictive correlations ranged from *r* = .82 to .93 corresponding to cross-validated R^2^ values between .67 and .87. Prediction errors were small (MAE = 0.28 to 0.45). All permutation tests showed ps < .001, confirming predictive performance significantly exceeded chance for every dimension. Predictive generalisation plots (Figure S2 in the supplemental materials) illustrate the close correspondence between observed and predicted values across folds, whilst null distribution histograms (Figure S3 in the supplemental materials) demonstrated that the observed predictive correlations lie well beyond chance distributions. Together, these results show strong evidence that the Barraclough & Wightman’s (2026) 5-dimensional model captures a stable, generalizable structure in how actions are perceived.

**Table 4.** Predictive performance.

| Rating dimension | Predictive $r$ | Predictive $R^2$ | MAE | $p$ |
| --- | --- | --- | --- | --- |
| Formidableness | .93 | .87 | .28 | .0001 |
| Friendliness | .92 | .85 | .29 | .0001 |
| Locomotion | .87 | .76 | .37 | .0001 |
| Abduction | .82 | .67 | .45 | .0001 |
| Environmental interaction | .84 | .72 | .41 | .0001 |

Note: All results based on 10-fold cross-validated regression and permutation tests with 1000 iterations.

We found a similar close relationship between factor scores and ratings for the original 4-factor model (Vinton et al., 2023, reported fully in the supplemental materials). Reliability analyses (Table S5), RDMs (Figure S6), correlation matrixes (Figure S7), tests of convergent and discriminant validity (Tables S8 & S9) and predictive performance (Table S10) calculations demonstrate that this earlier version of the 5D action space model can also systematically explain human action perception.

## Experiment 2 – metric scaling

### Methods

#### Participants

Thirty-four participants (33 female, 1 male; average age = 18.88, S.D. = .73) completed the rating experiment. This ensured we had a minimum of 30 participants; although no *a priori* power analysis was appropriate, we selected this number to satisfy the central limit theorem. Participants were recruited via the internal University of York SONA system and compensated with course credit; all reported normal or corrected-to-normal vision. This experiment was approved by the ethics committee of the Department of Psychology, University of York, and was performed in accordance with the ethical standards laid down in the 1964 Declaration of Helsinki.

#### Stimuli

We selected 8 pairs of actions to sample extensively 5D action space, and so linear interpolation in action space between action pairs varied predominantly along either one action space dimension, or combinations of dimensions (see Table 5). To morph between each action pair, we used the same procedure used by de la Rosa et al. (2016); Fedorov et al. (2018); Ferstl et al. (2017). We calculated the weighted average of the local joint angles of the two source actions (e.g. unscrewing and throwing). The weights of the contributing actions were set to generate 5 different morphs with equally spaced weights on the continuum between the two source actions. The 5 resulting morph actions were 0%, 25%, 50%, 75% and 100% of the second action. In the example here, morphs would thus range between 100% unscrewing + 0% throwing (0% morph) through to 0% unscrewing + 100% throwing (100% morph).

**Table 5.** Actions selected for morphing.

| Action 1 | Action 2 | Dimensions on which actions<br>*predominantly vary* |
| --- | --- | --- |
| Celebrating | Holding an aching back | Formidableness, Abduction |
| Ducking down | Applying cream to face | Locomotion |
| Pushing an object | Destroying paper | Formidableness |
| Stealing an object | Laughing | Friendliness |
| Supporting someone | Passing a football | Abduction |
| Unscrewing a bottle | Smashing an object | Friendliness, Locomotion Abduction |
| Unscrewing a bottle | Throwing a ball | Abduction |
| Trying to warm up | Mopping | Environmental interaction |

Morphing was conducted within the Unity 3D (Unity, San Francisco, CA. USA, https://unity.com) game engine by averaging between the joint angles recorded in the source action .bvh files. The resulting morphs were presented on screen by animating the same androgynous volumetric avatar as for Experiment 1. To generate action videos for the experiment, Unity 3D played back each of the action morphs in 25% steps between the 2 actions on screen (1280 × 1080 pixels, 60fps), and playback was recorded with OBS Studio (Bailey, 2017) and each action was saved as an .mp4 file. In total there were 40 different morphed actions with 5 morphs weights for the 8 different action pairs.

#### Procedure

All experiments were implemented within the online Gorilla platform (Anwyl-Irvine et al., 2021; Anwyl-Irvine et al., 2020). Once participants entered the site via an internet browser (Chrome, Firefox of Safari) on a laptop or desktop PC, participants completed a consent form, recorded their age and gender, and then instructions on the experimental task were displayed. The experiment consisted of 6 blocks of testing, during which participants were required to rate actions on Formidableness, Friendliness, Locomotion, Abduction, Environmental Interaction, and Intentionality; block order was randomised across participants. On each trial they first observed a fixation cross in the centre of a white screen for 500 ms, then a video of an action morph for its duration, and finally a response screen showing a 1-9 Likert scale where participants had to indicate their rating of the action by clicking an onscreen button with the mouse. Once a response was registered, the next trial commenced. If participants failed to respond within 2 s of the end of the action, a prompt “Please respond faster!” was shown on screen to encourage participants to indicate their immediate first impressions of the action. Each block contained 40 trials, during each one of the 40 different action morphs was shown, whilst action order was randomised within the block.

#### Analysis

Because of variance in the locations of source actions within the 5-dimensional action space, although morphed actions varied predominantly on one dimension, they also varied on the others simultaneously. To account for this, for each morph action we derived its model-based coordinates by linearly interpolating the factor scores of Actions 1 and 2 across the five morph weights (0, 25, 50, 75, 100%). This provided predicted factor scores (PredF) for all five action space dimensions (Formidableness, Friendliness, Locomotion, Abduction, Environmental interaction) for each of the 8 continua × 5 morph levels. We then fitted linear mixed-effects models (MATLAB, Statistics and Machine Learning Toolbox) to test whether participant ratings scaled with these model-derived factor scores. For each rating dimension, we regressed participants’ ratings on the corresponding predicted factor score (e.g., Formidableness ratings predicted from the model’s Formidableness coordinate for each morph). The models included random intercepts and random slopes for factor score by participant, to account for individual differences in both overall rating level and sensitivity to factor variation, as well as a random intercept for action continuum, to capture any baseline differences across continua. We compared linear models to quadratic alternatives to test whether a linear model was an optimal account of the relationship between factor scores and perceived action quality.

To test dimensional specificity, we asked whether ratings of each of the 5 action-quality dimensions (Dim1–Dim5) are predicted most strongly by the corresponding model-derived predictor (PredF1–PredF5), relative to the non-matching predictors. For each participant and rating dimension, we fitted an OLS regression:

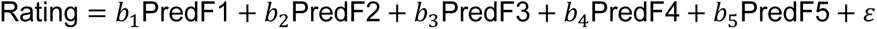

This generated a 5×5 matrix of slopes per subject, describing how strongly each PredF predicts each type of rating. The diagonal of this matrix corresponds to the dimension-matching effects (e.g., PredF1 → Dim1). The off-diagonals of the matrix correspond with nonspecific cross-predictions. Dimensional specificity predicts larger diagonal coefficients than off-diagonal coefficients, which we tested using paired t-tests.

### Results

#### Linearity

Across all five dimensions, perceived action qualities increased as the underlying morph weight moved from Action 1 (the action low in the quality) to Action 2 (the action high in the quality; see Figure 4). For Formidableness, morph % significantly predicted ratings of formidableness (β = .499, SE = .151, t = 3.29, p < .001), accounting for 21.9% of the variance. Comparing the linear model to a quadratic alternative revealed no improvement in fit (ΔAIC = .58; LRT p = .108), indicating approximately linearly scaling. For Friendliness, morph % significantly predicted ratings (β = .316, SE = .146, t = 2.17, p = .030), accounting for 23.7% of the variance in ratings. A quadratic model did not significantly improve fit (ΔAIC = –.75; LRT p = .264), indicating approximately linearly scaling. For Locomotion, morph % did not significantly predict ratings of Locomotion (β = .135, SE = .151, t = .89, p = .371), and a quadratic model did not significantly improve fit (ΔAIC = 1.39; LRT p = .159). For Abduction, the slope was positive but marginal (β = .313, SE = .168, t = 1.87, p = .062), and a quadratic model did not significantly improve fit (ΔAIC = 1.39; p = .066). This indicates that perceived changes in this quality scaled approximately linearly, albeit with weaker evidence compared to Formidableness, and Friendliness. Finally, for Environmental interaction, morph % significantly predicted ratings (β = 1.079, SE = .168, t = 6.42, p < .001), accounting for 37.9% of the variance in ratings. Comparing the linear model to a quadratic alternative revealed no improvement in model fit (ΔAIC = 1.81; LRT p = .051), indicating approximately linearly scaling. Together, these results show that the dimensions of Formidableness, Friendliness and Environmental interaction exhibit approximately linear perceptual scaling, supporting the interpretability of distances within the action-space model along these axes.

**Figure 4.**
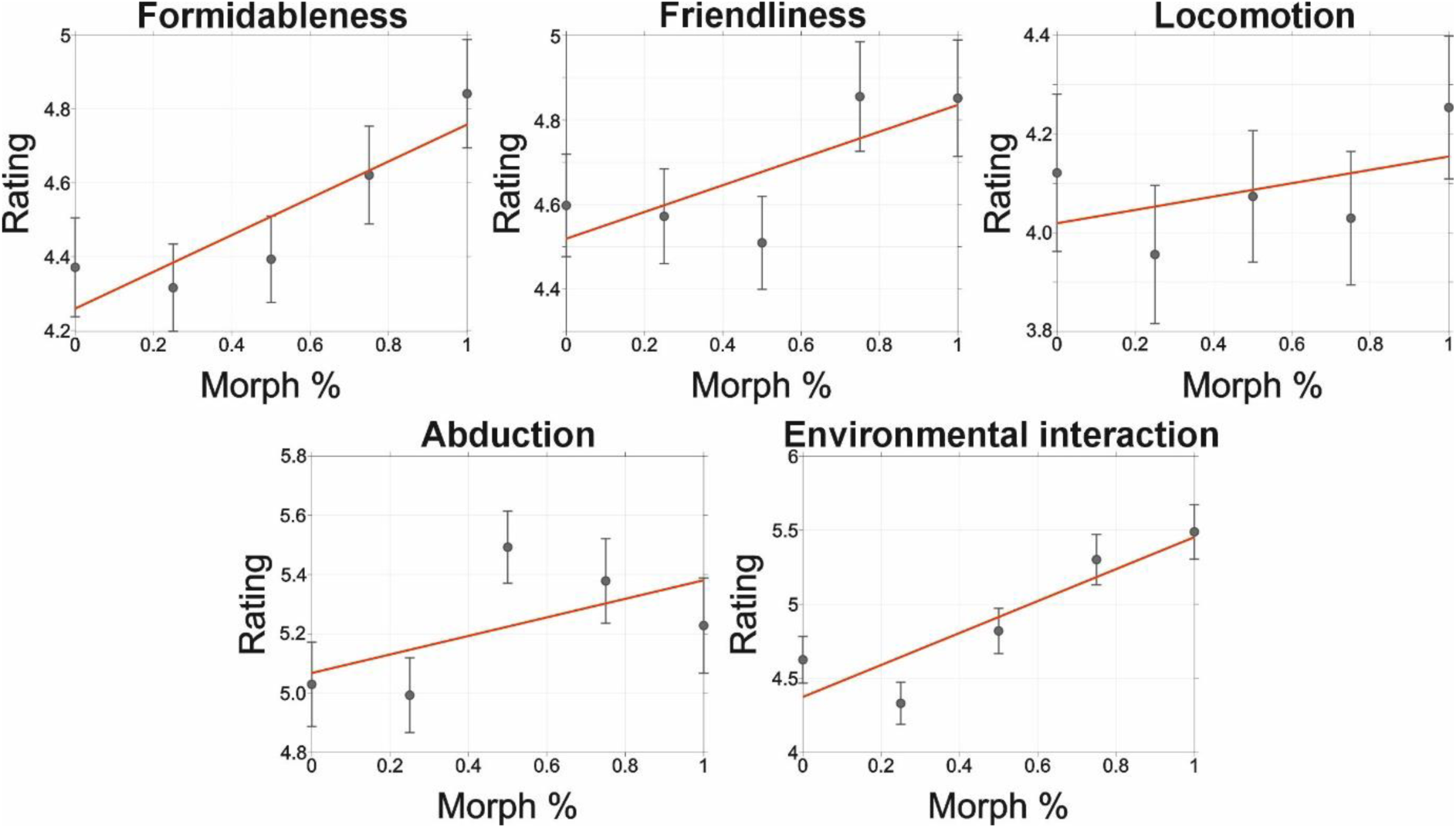
Participant ratings versus predicted factor values for ratings of the 5 different action quality dimensions Note: Grey: Markers – average participant ratings, error bars - standard error of the mean, Red: Linear mixed effect prediction between the factor score and participant ratings.

#### Dimension specificity

Dimension specificity was tested by comparing across participants on-diagonal coefficients against the average of the off-diagonal coefficients using Bayesian paired sample t-tests implemented via JASP (JASP-Team, 2021). Bayes factors indicated extreme evidence for our alternative hypothesis that on-diagonal coefficients were consistently larger indicating that each rating dimension was most strongly predicted by its corresponding PredF component (Formidableness: BF_10_ = 5.36×10^11^, error % = 4.65×10^-14^; Friendliness: BF_10_ = 9.53×10^14^, error % = 5.49×10^-17^; Locomotion: BF_10_ = 169,606, error % = 7.06×10^-11^; Abduction: BF_10_ = 4.21×10^8^, error % = 7.12×10^-13^, Environmental interaction: BF_10_ = 2.95×10^7^, error % = 8.07×10^-12^). These results provide clear evidence of dimension-specific tuning: that the predictor corresponding to each dimension contributes uniquely to explaining ratings along that dimension. This can be observed in the group-level beta matrix illustrating mean regression slopes across participants (Figure 5).

**Figure 5.**
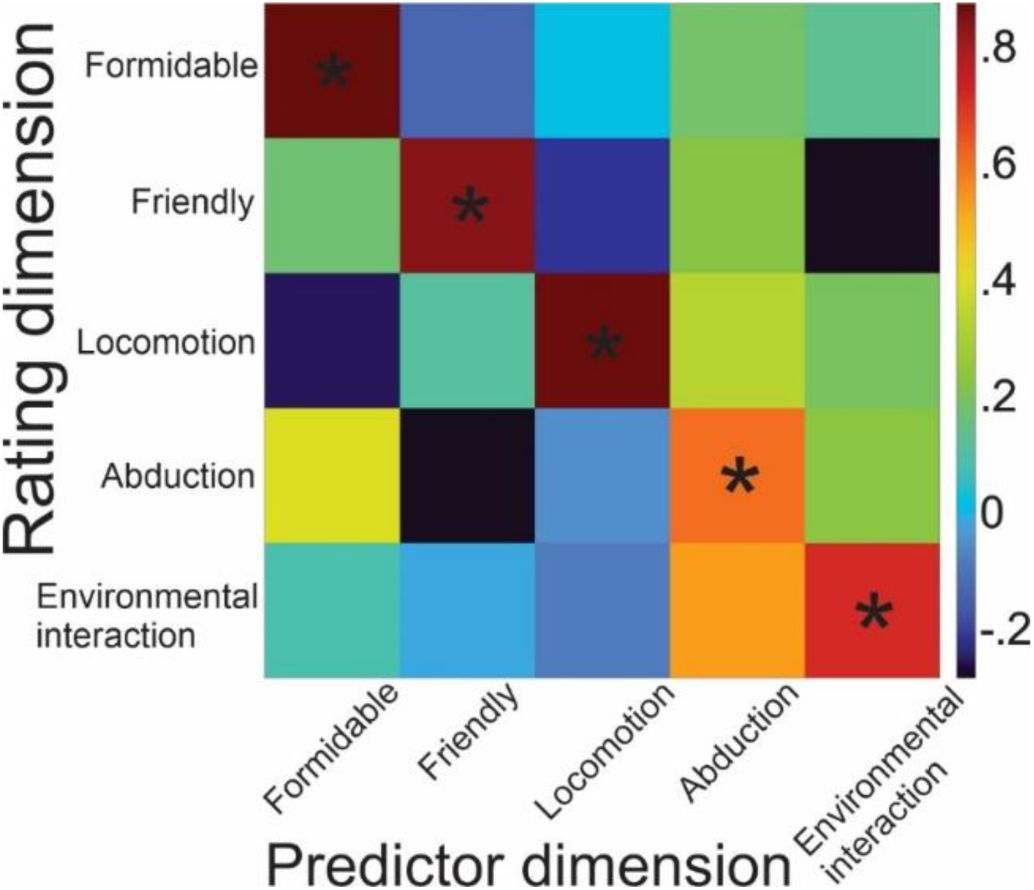
Group level beta matrix Note: Each cell denotes the mean regression coefficient relating a given rating dimension to a given PredF predictor. Rows – rating dimension (Dim1-Dim5); columns model predictors (PredF1–PredF5). The on-diagonal values are consistently largest in their respective rows, reflecting strong dimension-matching effects. Asterisks on the diagonal cells indicate dimensions where there was extreme evidence that the diagonal coefficients were larger than the mean off-diagonal coefficients.

To enable a comparison or these results with an equivalent analysis of the original 4-factor model (Vinton et al., 2023), we repeated these calculations with ratings of the 4 factors (Formidableness, Friendliness, Intentionality, Abduction) compared against predictions from the 4D model. Although intentionality ratings scaled with predicted factor score, this was not a linear relationship (Figure S11). However, the earlier 4D model also similarly and clearly demonstrated dimension specific tuning (Figure S12).

## Experiment 3 – computational dissociation analysis

### Methods

To determine the extent to which the 5D action space represents higher-order social-cognitive abstractions about actions rather than low-level physical regularities or linguistic labels, we compared the hypothesized model against both a structural kinematic baseline and a semantic representational space using RSA.

#### Kinematic baseline extraction

Structural information about each action was extracted directly from the raw .bvh motion-capture files on which all 240 actions were based (Vinton et al., 2023). To create a kinematic profile for each action, we extracted the raw motion-capture data, which included the hierarchical 3D rotations and translations for every joint across all frames of the action. To calculate a kinematic baseline for comparison, we represented each action using a postural-dynamic descriptor derived from these raw skeletal channels. This approach was chosen to capture the two fundamental components of action motion perception: the mean pose (the mean value of all skeletal channels across the duration of the action) and the global motion dynamics (represented by the temporal variance/standard deviation; Giese & Poggio, 2003). Unlike deep-learning architectures (e.g., Spatial Temporal Graph Convolutional Networks; ST-GCNs; Jiang et al., 2022; Yan et al., 2018) which may incorporate non-linear features optimized for action classification, this statistical kinematic baseline provides an interpretable ‘first-order’ representation of the physical signal. This allows for a clear dissociation between the raw postures and kinematics of the actions and the higher-order social-cognitive inferences captured by the 5D action space model. These descriptors were concatenated into a single high-dimensional structural feature vector for each action, providing a baseline of the raw physical information available in the action.

#### Semantic space construction

To model the semantic representation of the action stimuli, we constructed a representational space based on free-response labels to all 240 actions provided by 35 naive observers (data from Vinton et al., 2023). For each of the actions, all participant responses were first concatenated, and then a pre-trained Sentence-Transformer model (all-MiniLM-L6-v2) was used to map each set of descriptions into a 384-dimensional vector space in python 3.12.8. Unlike simple word-overlap models, this transformer-based approach (SBERT; Devlin et al., 2019) uses a bi-encoder architecture to capture distributional semantics, ensuring that conceptually related action labels (e.g., jumping and leaping) are positioned closely together even when they do not share identical vocabulary.

#### RDM construction and comparison

We constructed several RDMs (see Figure 6) to compare the geometry of the physical action space, the psychological 5D action space, and the semantic space. The kinematic structure RDM was generated by calculating the cosine distance between the structural feature vectors of every pair of actions. Cosine distance was selected to emphasize the relative pattern of joint movements rather than their absolute magnitude. The Semantic RDM was constructed similarly with cosine distance used to represent conceptual dissimilarity. Model RDMs consisted of six separate matrices based on the factor scores derived from the 5D action space model. These included a Full 5D Model RDM (using all 5D coordinates) and five unidimensional RDMs (e.g., a ‘Formidableness only’ RDM). Distances in these psychological spaces were calculated using Euclidean distance, representing the perceived psychological distance between actions within the action space model’s geometry.

**Figure 6.**
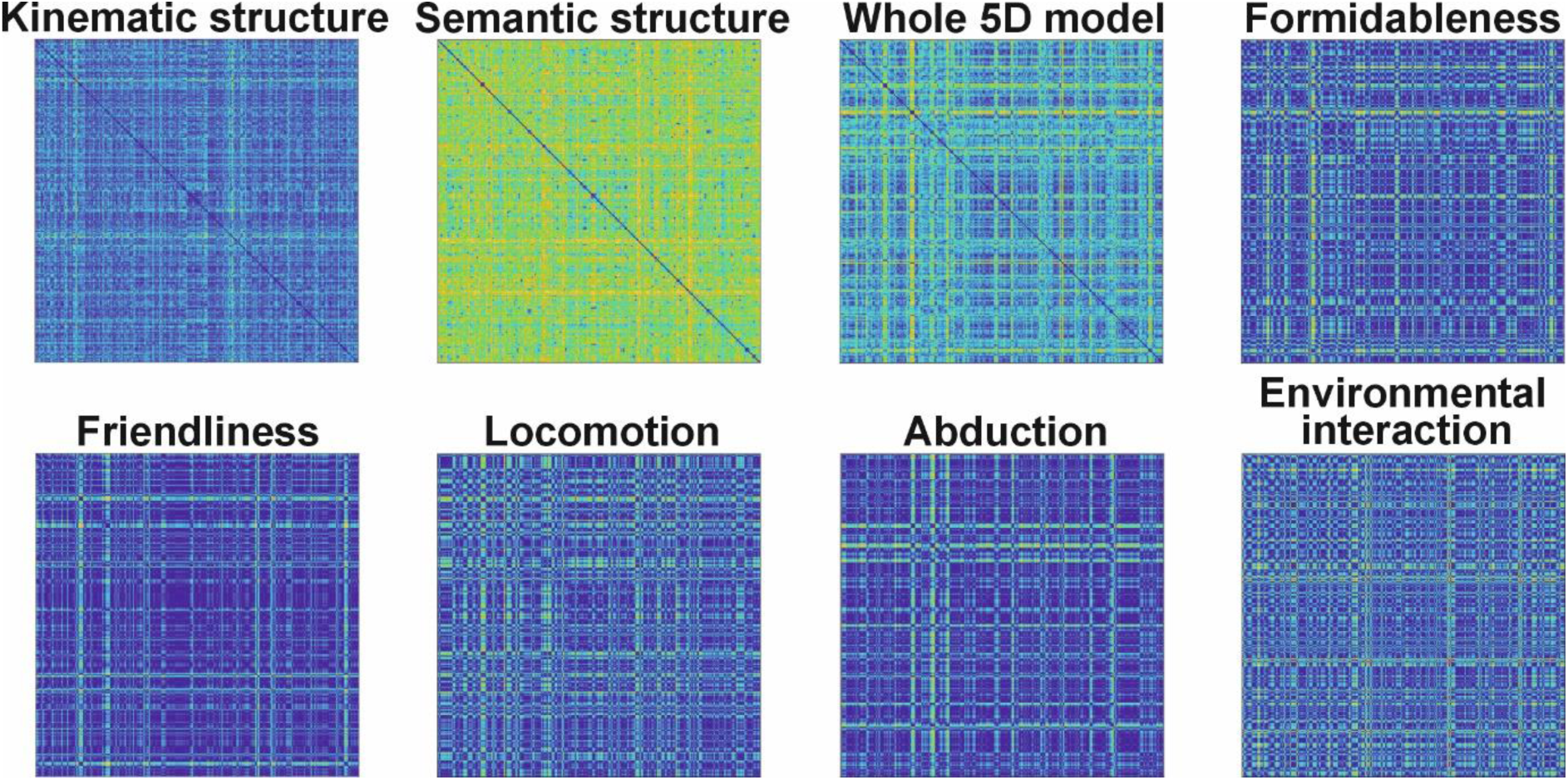
Representational dissimilarity matrices.

#### Statistical Validation and Multiple Regression RSA

The relationship between these spaces was initially assessed using Spearman’s rank correlations between the upper-triangular elements of the kinematic and semantic RDMs and each of the model RDMs. To determine if the 5D model captured unique semantic information not present in the raw physical motion, we conducted a Multiple Regression RSA. The Semantic RDM was treated as the dependent variable, with the Kinematic and 5D Model RDMs serving as standardized predictors. This allowed for the calculation of Beta weights (β) to determine the relative importance of physical vs. psychological information in predicting action meaning. Finally, we performed a partial correlation to assess the relationship between the 5D model and semantics while mathematically controlling for the influence of kinematics. This control tested whether any observed alignment between the 5D model and semantics is not merely a byproduct of low-level physical similarities.

### Results

While all correlations were highly significant (all ps < .00001, reflecting the high power of the 240 x 240 pairwise comparison matrices), the results revealed a clear hierarchical distinction between physical, psychological, and semantic layers of action representation. The kinematic baseline accounted for only a small proportion of the representational variance in the 5D model, with structural kinematics explaining just 1.19% of the shared rank order variance of the whole 5D model (ρ = .109). In contrast, the 5D model demonstrated a substantially stronger alignment with the semantic representational space (ρ = .370; ρ^2^ = 13.69%) accounting for approximately 11 times more variance in semantic labels than in raw joint kinematics. This suggests that the geometry of the 5D action space is primarily organized around more conceptual properties of the actions rather than lower-level structural information.

Dimension-specific analyses further highlighted this dissociation (Figure 7). When compared against the kinematic baseline, physical kinematics explained variance in the Locomotion dimension the most (ρ = .142, ρ^2^ = 2.02%) but less than 0.2% of the variance in the more social-evaluative dimensions such as Friendliness and Formidableness (both ρ = .036). However, when compared against the semantic space, all dimensions showed closer alignment, and with a different profile. The Environment interaction (ρ = .258; ρ^2^ = 6.66%) and Friendliness (ρ = .227; ρ^2^ = 5.15%) dimensions showed the strongest semantic relationship with the 5D model, with Locomotion showing the least (ρ = .165; ρ^2^ = 2.72%).

**Figure 7.**
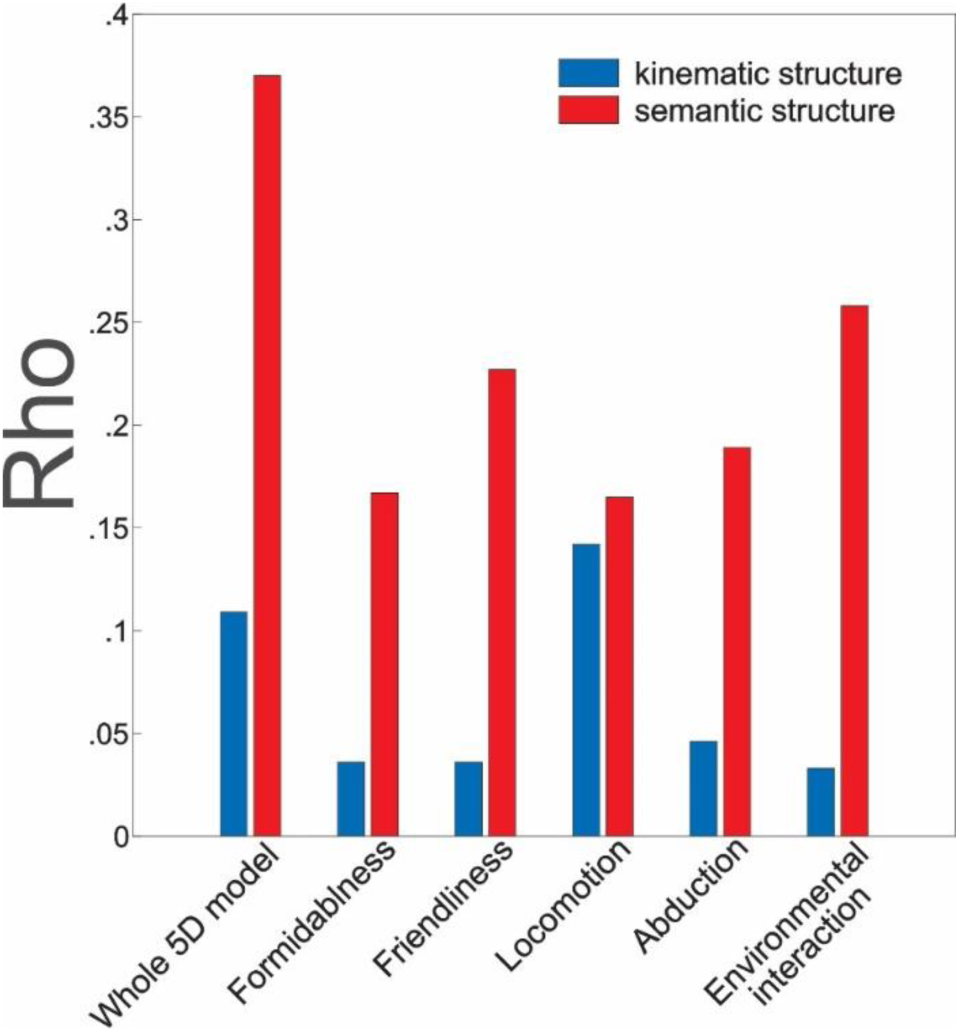
Spearman rank correlations between kinematic and semantic structures and the action space model and its individual dimensions

To confirm that the action space model captured unique cognitive information not present in the physics of the actions, we conducted a multiple regression RSA. The combined model (Kinematics + 5D Model) explained 17% of the total variance in action semantics (R^2^ = .170). Critically, standardised regression coefficients revealed that the 5D model (β = .374) was a significantly stronger predictor of action meaning than raw kinematics (β = .136). This interpretation was further supported by a partial correlation analysis: when mathematically controlling for the influence of the structural kinematic RDM, the relationship between the 5D model and the semantic space remained almost entirely intact (Partial r_s_ = .364, p < .001 compared to ρ = .370). These results demonstrate that the 5D action space predominantly captures more abstract, social inferences that are not only irreducible to first-order skeletal statistics but also provide a framework for mapping physical human actions onto distal social meaning. Finally, a similar profile of results was found for Vinton et al.’s (2023) 4D model of action space (supplemental materials Figure S13), demonstrating the structural and conceptual equivalency of these two models.

## Discussion

We conducted three experiments to evaluate the psychological validity, metric scaling and quantify the hierarchical position within the social visual processing network of the 5D action space model identified by Barraclough and Wightman (2026). In the first experiment we demonstrated the psychological and geometric validity of the model, providing evidence to suggest it is a valid cognitive map. Clear methodological convergence showed that it reflects real-world human perception (Indow & Kanazawa, 1960; Wiggins, 1979) and can act as a universal, generative framework where continuous geometric coordinates systematically determine qualitative attributes (Edelman & Intrator, 2003).

A necessary first step toward this methodological convergence was demonstrating that the dimensions capture a stable consensus across independent observers rather than idiosyncratic rater noise (Wiggins, 1979). The intraclass correlation coefficients (ICC) and Cronbach’s alpha metrics for all five action qualities were very high, with the primary dimensions (Formidableness, Friendliness, Locomotion, Abduction and Environmental interaction) all exceeding .86. Even the highly context-dependent dimension of Environmental interaction demonstrated robust reliability (ICC = .862). These data indicate that human observers use a highly consistent, shared representation when evaluating actions on these dimensions.

Using Representational Similarity Analysis (RSA), we demonstrated a clear topological alignment between the latent factor space and behavioural rating space (*r* = .79). This finding directly parallels validation paradigms successfully used in other psychological domains. For example, Thornton and Tamir (2020) used RSA to establish a structural match between a 3D mental state space and behavioural ratings, while Stolier et al. (2018) and Jozwik et al. (2022) utilized similar geometric correspondences to show that the conceptual structure of trait spaces determines the perceptual representation of faces. Our RSA results extend this principle into the action domain. Within the frameworks of Gärdenfors (2004a, 2004b) and Shepard (1962a, 1962b), the distance between items within a true conceptual space map onto their psychological similarity. The strong correlation between our factor-derived and rating-derived representational dissimilarity matrices (RDMs) confirms that the 5D latent space accurately preserves this global geometry, demonstrating that its internal metrics faithfully mirror the distances on the different meaningful action quality dimensions that humans use to evaluate actions.

Because the original latent factor model was derived using an oblique rotation, it explicitly allowed the underlying factors to correlate, consequently there were moderate inter-factor associations (*r* = .04 to .56). Crucially, these correlations reflect ecological realities in actions rather than statistical redundancy. For example, ratings of Abduction correlated strongly with those of Formidableness (r = .70). In biological systems, this relationship is highly functional: expanding one’s posture (high abduction) is a universal physical strategy deployed across species to signal dominance and power (Altschul, 2024; Darwin, 2025). Importantly, our variance partitioning and multiple regression analyses showed the dimensions maintain clear discriminant validity, despite these real-world ecological overlaps. For every dimension, the matched latent factor was the dominant predictor of its corresponding behavioural rating (βs = .65 to .91), explaining large proportions of unique variance (up to R^2^ = .72 for *Friendliness*). This demonstrates that the 5D model respects the natural correlational structure of stimulus information while successfully isolating distinct, independent dimensions of social-cognitive inference.

Perhaps the most persuasive evidence for the psychological validity of the 5D action space comes from the 10-fold cross-validated regression analysis, providing a direct test of the model’s generative ability. Predictive performance was consistently excellent across all five rating dimensions, with cross-validated predictive correlations ranging from *r* = .82 to .93, accounting for up to 87% of the total variance in held-out data. This predictive power closely parallels generative modelling in other domains where an item’s coordinates reliably predict qualitative feature attributions in unobserved data (Jozwik et al., 2022; Thornton & Tamir, 2020), whilst our findings satisfy the criteria for computational models of visual systematicity (Edelman & Intrator, 2003). Human perception is thought to rely on graded, continuous representational spaces rather than discrete categories. By demonstrating that an action’s specific coordinates within the 5D space systematically and deterministically dictate its perceived qualities, we provide a formal empirical basis for this continuous dimensional action space. Rather than simply categorising actions, the human brain appears to use this continuous geometric space as a universal, generative framework to govern judgments about human actions.

The objective of Experiment 2 was to determine whether the 5D action space functions as a true perceptual metric. A defining characteristic of valid psychological spaces are that metric distances correspond to perceptual differences (Indow & Kanazawa, 1960; Shepard, 1987). By generating morph continua between exemplar actions, we tested whether geometric distances within the statistically defined action space selectively and linearly predicted the intensity of perceived action qualities. Our results provided clear evidence that the primary social-evaluative dimensions of this space behave as psychophysical scales. For the dimensions of Formidableness, Friendliness, and Environmental Interaction, we observed significant linear relationships between the underlying morph weights and participant ratings, in comparison to quadratic alternatives. This absence of non-linear effects suggests that these dimensions function as do not simply categorize actions; rather, they function as graded, continuous metrics where every unit of change in the model’s geometric coordinates corresponds to a proportional shift in subjective perception. These results are commensurate with evidence for metric validity in other complex biological domains (e.g. Jozwik et al., 2022; Troje, 2002). For example, with the gait space model proposed by Troje (2002), linear decompositions of human motion reliably predict the intensity of perceived social attributes. By demonstrating that our model-derived coordinates can be used to synthesize novel, intermediate actions that are perceived with predictable intensity, we provide a formal computational basis for the idea that human perception relies on a continuous, systematic map of the action domain (Edelman & Intrator, 2003).

Beyond simple linearity, valid multidimensional models require dimension-specific tuning - meaning that a change along one axis of the space should selectively alter the perception of that specific quality without affecting unrelated dimensions. Our Bayesian analysis provided extreme evidence that the on-diagonal regression coefficients were consistently larger than off-diagonal associations. This selective tuning indicates that the 5D action space possesses a high degree of discriminant metric validity. While social actions are often ecologically correlated (e.g., the relationship between abduction and formidableness), we demonstrate that the human social perceptual system can isolate the specific variance associated with each latent dimension. This correspondence between the model’s structure and perceptual tuning is a corollary of recent findings in the face domain (Jozwik et al., 2022), suggesting that the brain organizes complex social stimuli into selectively tuned representational channels to facilitate rapid social inference.

While the social-evaluative dimensions of Formidableness and Friendliness showed robust linear scaling, the results for Locomotion and Abduction were less pronounced. Locomotion did not significantly scale with morph percentage, and the evidence for Abduction was marginal. This suggests that human sensitivity to metric changes may not be uniform across all dimensions of the action space. One possibility is that Abduction might function more as a categorical anchor (e.g., ‘is the actor abducting or not?’ see Figure 4). Alternatively, the lack of significant scaling in these kinematic-heavy dimensions may be a byproduct of the morphing procedure itself; if the physical differences between the selected exemplars were not sufficiently distinct along those specific axes, the resulting perceptual gradients would naturally be flatter (e.g. as for Locomotion; see Figure 4).

Collectively, Experiment 2 confirms that the 5D action space model satisfies Shepard’s (1987) *Universal Law of Generalization*. Our results show that the geometric distance between two points in the 5D action space selectively and (largely) linearly predicts the qualitative differences humans perceive in the underlying actions. This demonstrates that the model is more than a statistical description; it is a perceptual metric that captures the systematicity of human action observation.

A central aim of this study was to quantify the hierarchical position of 5D action space model and its constituent dimensions within the social visual processing network. By comparing the geometry of the model against both a structural kinematic space and a high-dimensional semantic space, we identified a clear dissociation between the physics of human actions and their social-cognitive representation. The dissociation between the 5D action space and the kinematic space cannot be attributed to a lack of inter-rater reliability in the development of the 5D model. There were high levels of agreement across observers for the constituent characteristics of each model factor (mean kappa = .808; Barraclough & Wightman, 2026). Given that Kappa is a conservative metric that corrects for chance agreement, this demonstrates that the consensus ratings used in the model were highly stable. These values effectively serve as an empirical noise ceiling; indicating that while humans agree on the meaning of these actions, that meaning is largely absent from the raw physical signal (ρ^2^ = .0119).

The most striking finding was the disparity in variance explained by kinematics compared with semantics. While the structural descriptors captured joint-rotation information and motion energy available in the actions, they accounted for less than 0.2% of the variance in the principal model dimensions of Friendliness and Formidableness. In contrast the 5D model demonstrated a much more robust alignment with the semantic space (ρ = .370). Indeed, the model was approximately 11 times more effective at predicting this abstract representation of actions than their kinematic properties. This suggests that the meaning of actions captured by the 5D model is less a property of action kinematics, rather the result of more top-down social inferences (de la Rosa et al., 2014; Fedorov et al., 2018). This greater alignment with semantics suggests that Barraclough & Wightman’s (2025) 5D model of action space can bridge the “Social-Cognitive Gap” observed in current AI architectures (Garcia et al., 2025; Garcia et al., 2024), where even the best models of actions like ST-GCNs (Yan et al., 2018) fail to match human evaluations of actions because they lack a dedicated social-cognitive viewpoint.

The multiple regression RSA provided the most direct evidence of the model’s unique cognitive utility. Standardized regression coefficients revealed that the 5D model (β = .374) was a substantially more powerful predictor of action meaning than raw kinematics (β = .136). Crucially, the partial correlation between the model and semantics remained nearly identical even after mathematically controlling for all kinematic similarities. This indicates that the 5D action space is not just a representation of action movement; it captures a unique layer of semantic variance that is independent of the raw skeletal statistics.

Critically, our results allow the evaluation of the hierarchical position of the different dimensions within the social processing network, as well as a validation of our RSA methodology. Locomotion showed the highest correlation with structural kinematics (ρ^2^ = .0202), explaining significantly more variance than any other dimension. Conceptually, this shows that our RSA was sensitive enough to detect physical regularities in the actions when they existed, as Locomotion is defined by the physical movement of the body through space, its representation is more closely tied to kinematics. Next closest linked to kinematics were the dimensions of Formidableness and then Abduction. These dimensions are associated with each other and both involve the representation of the speed and power of body movements and information on whether the body parts are moving towards or away from the body (Barraclough & Wightman, 2026; Vinton et al., 2023). In contrast, the dimensions of Friendliness and Environmental interaction remained dissociated from kinematics, yet strongly tied to semantics, demonstrating that these dimensions are fundamentally different in their computational origin. Friendliness is a social-evaluative dimension, and our results suggest that there is more than one way to convey friendliness. Furthermore, evaluating action friendliness requires inferences about the mental state of the actor, and consequently requires more abstract processing in mentalising system regions (Saxe & Kanwisher, 2003; Spunt et al., 2011). The reason why Environmental interaction is most related to a semantic space is two-fold. First, the actions are motion-capture animated avatars where objects and the physical environment are not visible. During observation we must infer the presence of objects from observing the actions. This process involves accessing conceptual knowledge of the actions at more abstract levels of processing rather than the physical constraints of the objects on which they act (Chen et al., 2023; Wurm et al., 2015). Second, this dimension might reflect the evaluation of more than whether actions involve interactions with the physical environment or not. Although not originally classified as substantive, whether the actions are person-directed or not also loaded negatively onto this dimension (Barraclough & Wightman, 2026). Consequently, this dimension might reflect a somewhat more complex continuum between actions that are directly related towards objects/physical environment through to actions that are directed towards other individuals (cf. Tarhan & Konkle, 2020; Wurm et al., 2017). Deriving this information from these actions would thus require access to more conceptual action representations (Tucciarelli et al., 2019; Wurm & Caramazza, 2019, 2021).

The low overall correlation with kinematics contrasted with the robust semantic alignment suggests the 5D model represents a highly abstracted cognitive representation. By using volumetric, featureless avatars, this effectively isolates the ‘pure’ action (posture and kinematics) from other potentially confounding variables like facial expressions, body shape, clothing etc. Even in this isolated state, the human brain appears to discard most raw kinematic data in favour of a low-dimensional representation optimised for social utility. This finding supports a hierarchical view of the action observation network (AON; Decety & Grezes, 1999). While the early visual cortex, and potentially the superior temporal sulcus (STS), may track the physical “structure” of motion (Giese & Poggio, 2003), the representational space tested here likely resides at later stages of processing - where posture and kinematics are transformed into meaning.

In conclusion, here we establish that the 5D action space model is a perceptually scaled, continuous geometric framework that reliably maps how the human brain transforms action kinematic information into social meaning optimised for social utility. Whist the dimension of Locomotion is most related to action kinematics, the abstract dimensions of Friendliness and Environmental interaction instead capture more abstract, inferential information of human intent and goal-directed behaviour. By delineating this higher-order layer of action representation, the 5D action space model provides a robust empirical basis for understanding the systematic way humans evaluate and organize the complexities of social actions.

## Supplemental materials

## Experiment 1 – testing the perceptual validity of the 5D model

**Table S1.** Full regression results and correlations for each 4D model factor predicting each rating.

| Factor score | Formidableness rating | Friendliness rating | Locomotion rating | Abduction rating | Environmental interaction rating |
| --- | --- | --- | --- | --- | --- |
| Formidableness | $\beta = .678, R^2 = .273$ | $\beta = -.092, R^2 = .005$ | $\beta = -.065, R^2 = .003$ | $\beta = .026, R^2 = .000$ | $\beta = .220, R^2 = .029$ |
| Friendliness | $\beta = -.255, R^2 = .056$ | $\beta = .912, R^2 = .718$ | $\beta = -.045, R^2 = .002$ | $\beta = .006, R^2 = .000$ | $\beta = -.077, R^2 = .005$ |
| Locomotion | $\beta = .180, R^2 = .026$ | $\beta = .038, R^2 = .001$ | $\beta = .835, R^2 = .563$ | $\beta = .145, R^2 = .017$ | $\beta = .218, R^2 = .038$ |
| Abduction | $\beta = .212, R^2 = .027$ | $\beta = -.115, R^2 = .008$ | $\beta = .164, R^2 = .016$ | $\beta = .740, R^2 = .330$ | $\beta = .250, R^2 = .038$ |
| Environmental interaction | $\beta = .237, R^2 = .056$ | $\beta = -.167, R^2 = .028$ | $\beta = -.052, R^2 = .003$ | $\beta = .065, R^2 = .004$ | $\beta = .645, R^2 = .413$ |

**Note:** Standardized regression coefficients ( $\beta$ ) and unique variance explained ( $R^2$ ) are shown. Bold entries indicate the ‘matched’ factor–rating pairings. Together, these results demonstrate both convergent and discriminant validity of the 5D action space dimensions, supporting their interpretation as distinct but related qualities of actions.

**Figure S2.**
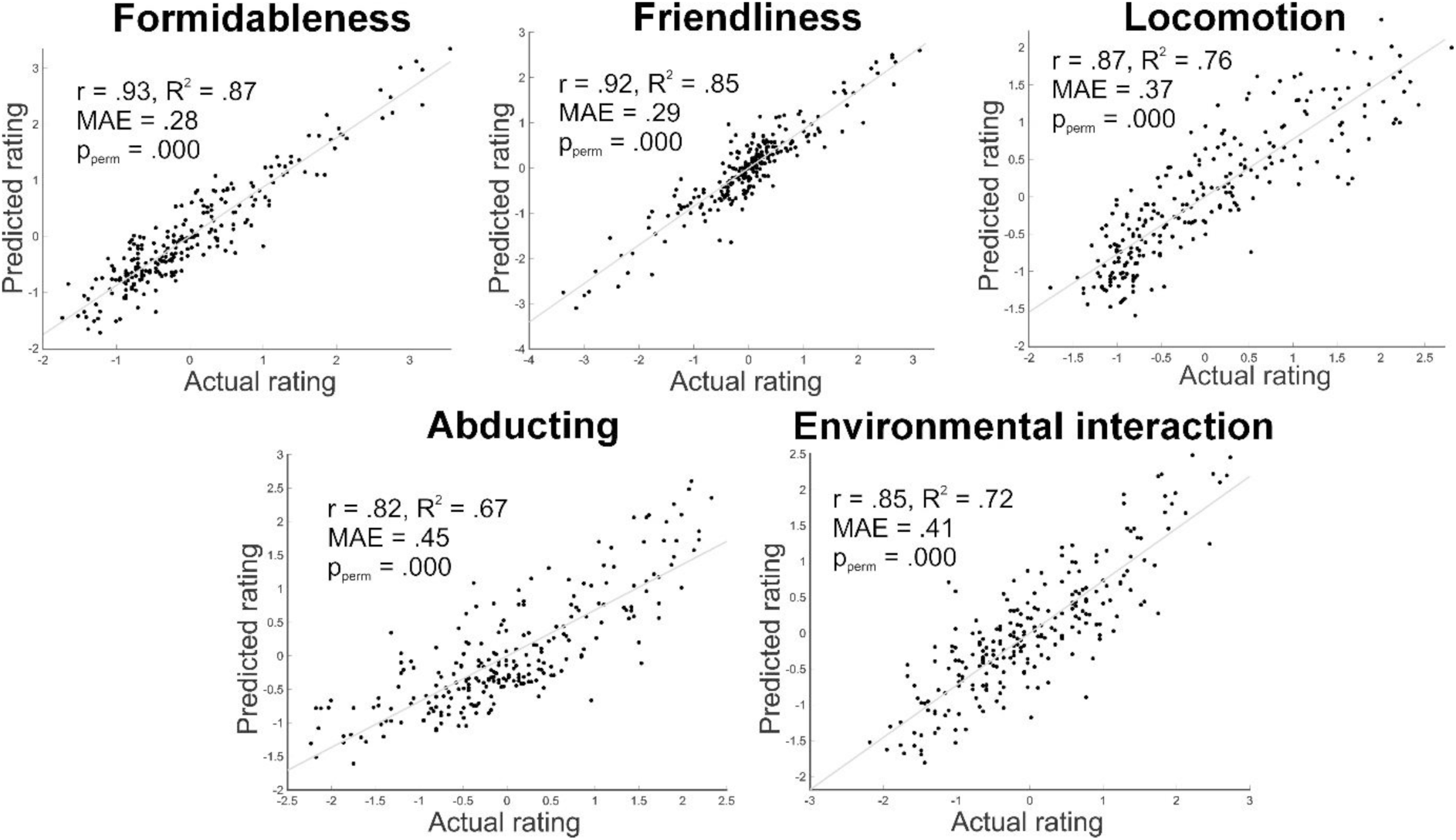
Predictive generalisation Note: Plots showing the relationship between predicted ratings and actual ratings for all five action space dimensions. Values for the Pearson correlation coefficient (r), goodness of fit (R^2^), Mean Absolute Error (MAE), and p value for the permutation analysis (pperm) are shown.

**Figure S3.**
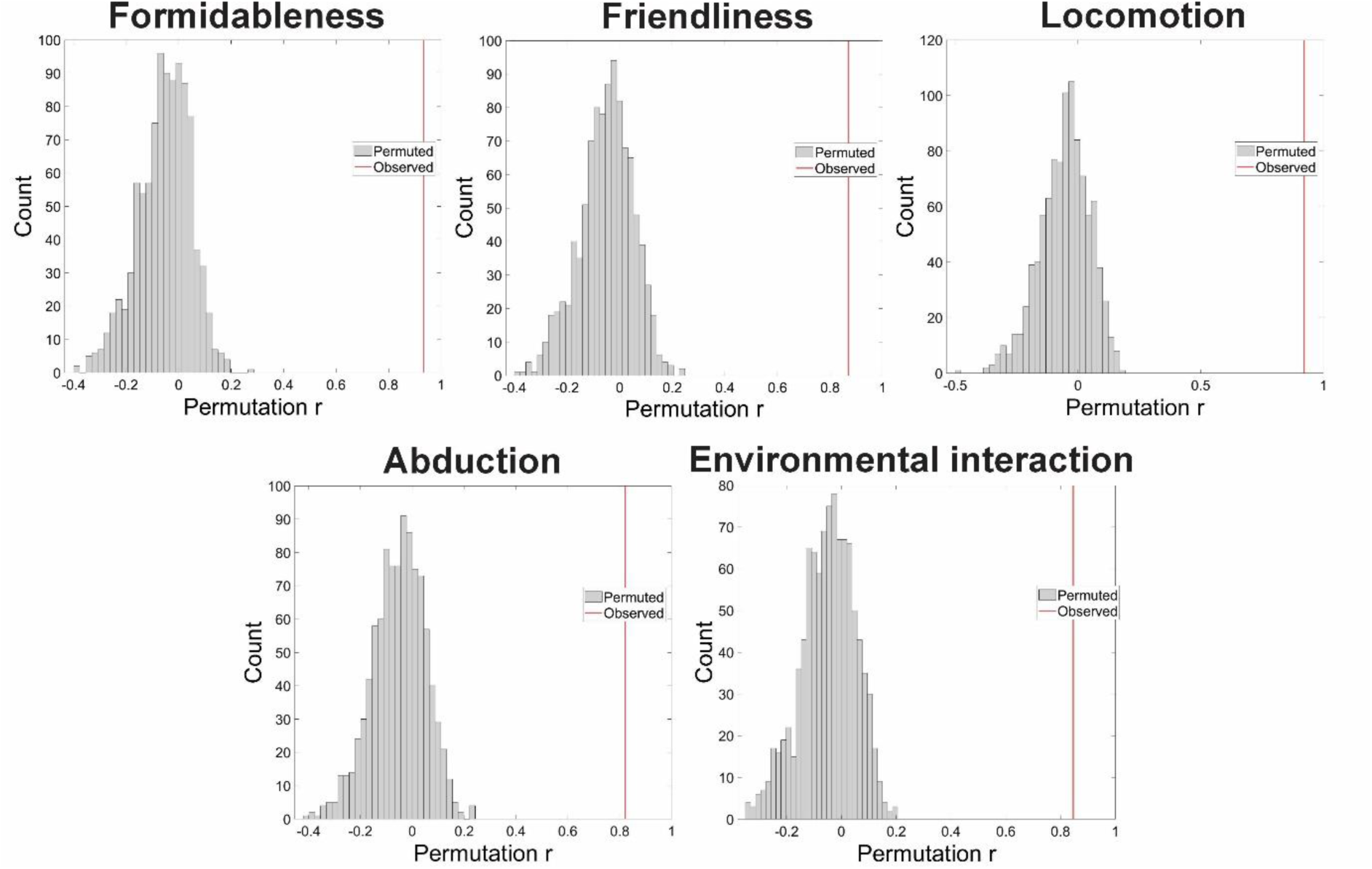
Null distributions from permutation tests Note: For all 5 dimensions of action space, observed predictive correlations lie far beyond the empirical chance distributions.

## Experiment 1 – testing the perceptual validity of the 4D model

**Table S4.**
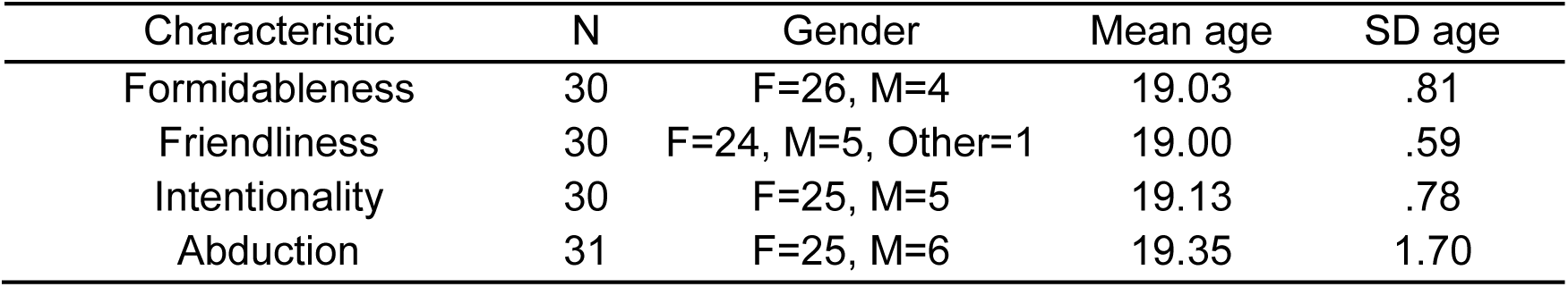
Participant demographic information.

| Characteristic | N | Gender | Mean age | SD age |
| --- | --- | --- | --- | --- |
| Formidableness | 30 | F=26, M=4 | 19.03 | .81 |
| Friendliness | 30 | F=24, M=5, Other=1 | 19.00 | .59 |
| Intentionality | 30 | F=25, M=5 | 19.13 | .78 |
| Abduction | 31 | F=25, M=6 | 19.35 | 1.70 |

For those rating actions on intentionality, the instructions were to: “Rate the actions on how intentional they appear on a scale from 1 to 9. With 1 = Very unintentional, 5 = Neither unintentional nor intentional, 9 = Very intentional. For example, bouncing a basketball might be seen as very intentional, tripping and falling to the floor might be seen as very unintentional, whilst playing on a phone might be seen as neither unintentional nor intentional.”

**Table S5.** Interclass correlation coefficients (ICC) and Cronbach’s alpha for all 4 action qualities.

| Characteristic | ICC<br>average<br>measure | 95% CIs | F-statistic | Interpretation | Cronbach’s<br>alpha |
| --- | --- | --- | --- | --- | --- |
| Formidableness | .950 | (.941, .959) | $F(239,6931) = 20.07, p < .001$ | Excellent | .95 |
| Friendliness | .953 | (.944, .961) | $F(239,6931) = 21.33, p < .001$ | Excellent | .95 |
| Intentionality | .947 | (.937, .957) | $F(239,6931) = 19.02, p < .001$ | Excellent | .95 |
| Abduction | .937 | (.925, .948) | $F(239,7170) = 15.87, p < .001$ | Excellent | .94 |

**Figure S6.**
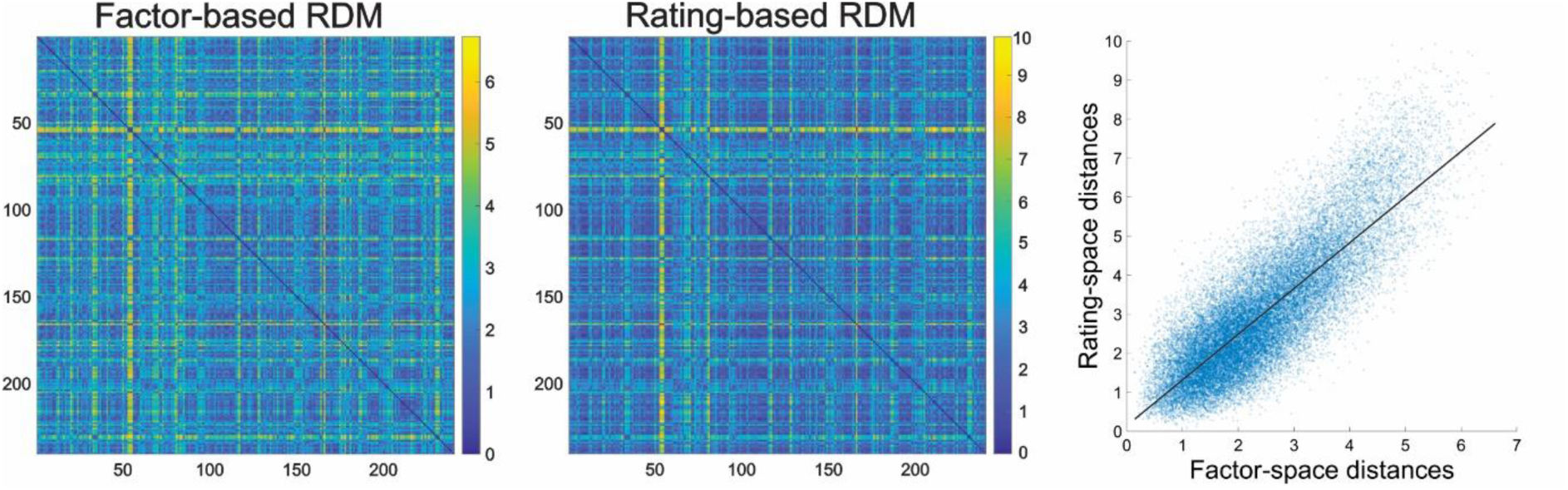
Representational dissimilarity matrices for 4D model Note: The left panel illustrates the dissimilarity matrix for actions derived from CFA-EFA factor scores (Vinton, Preston et al. 2023). The middle panel illustrates the dissimilarity matrix for actions derived from ratings of actions against 4 different qualities. The right panel illustrates the relationship (Spearman’s r=.83, p=.0001) between distances between pairs of actions (each marker) in rating space and factor score space; the black line indicates the best-fit regression line. A Mantel test indicated that the geometry of the 4D factor space predicted judgments of action qualities (Spearman’s r = .80, p=.0001).

**Figure S7.**
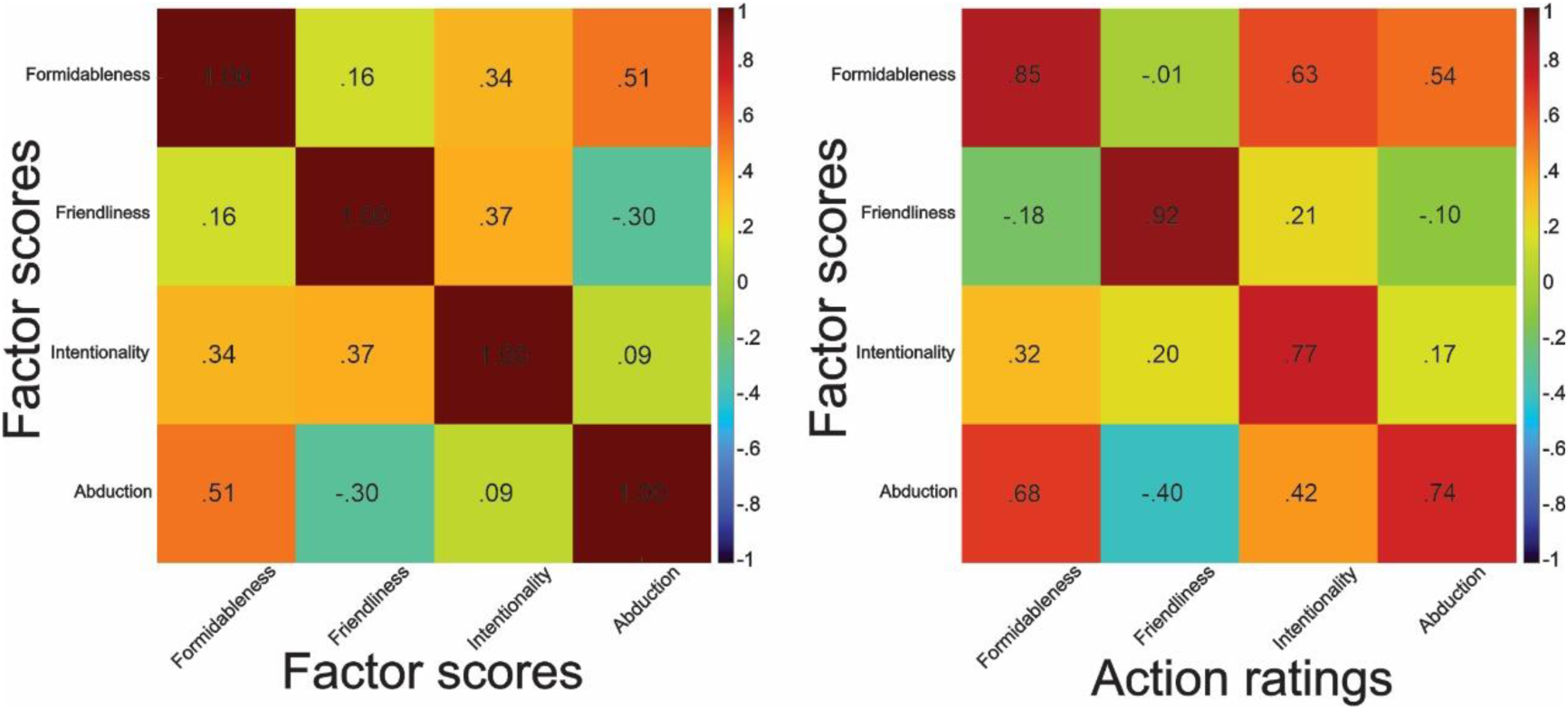
Action quality correlation matrices for 4D model Note: Left panel illustrates the factor score correlation matrix; inter-factor correlations were moderate in size (rs .16 to .51). Right panel illustrates action ratings correlated with factor scores and were strong (rs .74 to .92).

**Table S8.** Regression and variance partitioning for ratings predicted by 4D model factors.

| Rating (DV) | Target factor (IV) | b (target) | Full model $R^2$ | Unique $R^2$ (target) | Largest cross-predictor b | Shared $R^2$ | VIF |
| --- | --- | --- | --- | --- | --- | --- | --- |
| Formidableness | Formidableness | .74 | .87 | .33 | Abduction: $\beta = .31$ | .43 | 1.66 |
| Friendliness | Friendliness | .96 | .88 | .64 | Formidableness: $\beta = -.30$ | .22 | 1.43 |
| Intentionality | Intentionality | .66 | .79 | .34 | Formidableness: $\beta = .31$ | .37 | 1.23 |
| Abduction | Abduction | .66 | .58 | .25 | Formidableness: $\beta = .19$ | .30 | 1.69 |

**Table S9.** Full regression results and correlations for each 4D model factor predicting each rating.

| Factor score | Formidableness rating | Friendliness rating | Intentionality rating | Abduction rating |
| --- | --- | --- | --- | --- |
| Formidableness | $\beta = .745, R^2 = .335$ | $\beta = -.081, R^2 = .004$ | $\beta = .306, R^2 = .056$ | $\beta = .187, R^2 = .021$ |
| Friendliness | $\beta = -.301, R^2 = .064$ | $\beta = .956, R^2 = .640$ | $\beta = -.014, R^2 = .000$ | $\beta = .057, R^2 = .002$ |
| Intentionality | $\beta = .162, R^2 = .021$ | $\beta = -.120, R^2 = .011$ | $\beta = .657, R^2 = .338$ | $\beta = .024, R^2 = .000$ |
| Abduction | $\beta = .193, R^2 = .022$ | $\beta = -.065, R^2 = .003$ | $\beta = .207, R^2 = .025$ | $\beta = .655, R^2 = .254$ |

**Note:** Standardized regression coefficients ( $\beta$ ) and unique variance explained ( $R^2$ ) are shown. Bold entries indicate the ‘matched’ factor–rating pairings. Together, these results demonstrate both convergent and discriminant validity of the 4D action space dimensions, supporting their interpretation as distinct but related qualities of actions.

**Table S10.** 4D model predictive performance.

| Rating dimension | Predictive $r$ | Predictive $R^2$ | MAE | $p$ |
| --- | --- | --- | --- | --- |
| Formidableness | .93 | .86 | .30 | .0001 |
| Friendliness | .94 | .87 | .27 | .0001 |
| Intentionality | .88 | .78 | .36 | .0001 |
| Abduction | .75 | .56 | .51 | .0001 |

Note: All results based on 10-fold cross-validated regression and permutation tests with 1000 iterations

## Experiment 2 – testing linear scaling in the 4D model

To evaluate the 4-dimensional model we additionally tested how perceived action intentionality increased as the underlying morph weight moved from Action 1 to Action 2 (this relationship for formidableness, friendliness and abduction were tested for the 5D model and are reported in the main paper). For intentionality, the effect of morph % was strong (β = 1.306, SE = .179, t = 7.32, p = 4.4 × 10⁻¹³), explaining 27.7% of the variance. Unlike the other dimensions, however, the quadratic term significantly improved the fit over the linear model (ΔAIC = 3.97; p = .0146; Figure Y). This indicated a systematic non-linearity in how observers perceived progression in intentionality. That is, equal physical steps in the model coordinate did not correspond to equal perceptual steps: observers vary in their sensitivity in different regions along this dimension, suggesting that this dimension may compress or expand perceptual differences in a manner not captured by a strictly linear representation (see Figure S11).

**Figure S11.**
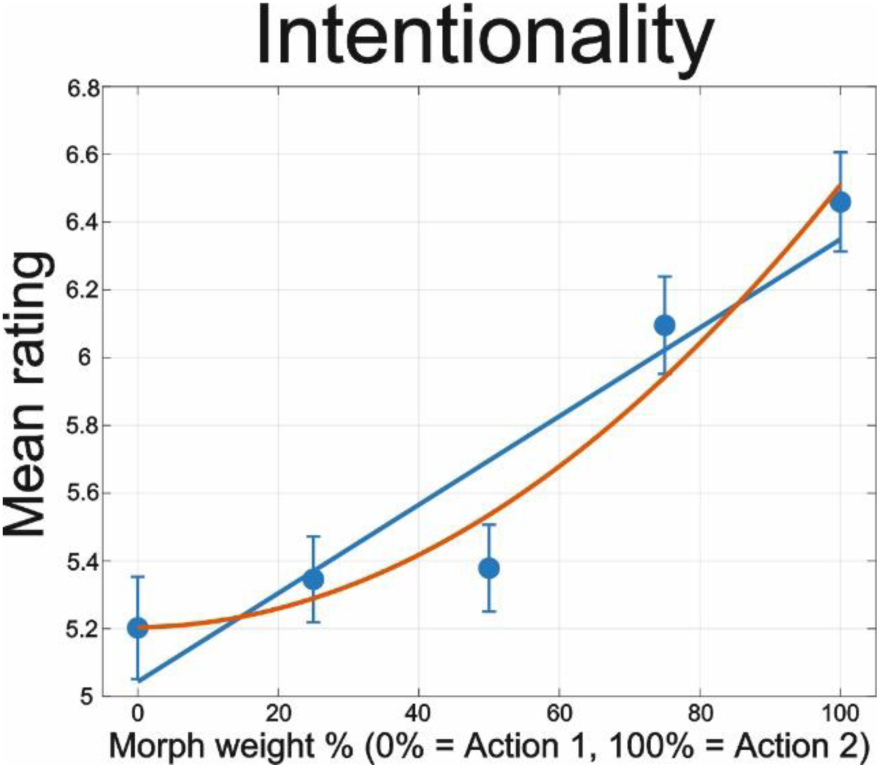
Participant ratings versus predicted factor values for intentionality from the 4D model Note: Blue: Markers – average participant ratings, error bars - standard error of the mean, linear mixed effect prediction between the factor score and participant ratings. Red line: quadratic mixed effect prediction.

Dimension specificity was tested by comparing across participants on-diagonal coefficients against the average of the off-diagonal coefficients (see Figure S12) using Bayesian paired sample t-tests implemented via JASP (JASP-Team, 2021). Bayes factors indicated extreme evidence for our alternative hypothesis that on-diagonal coefficients were consistently larger indicating that each rating dimension was most strongly predicted by its corresponding PredF component (formidableness: BF_10_ = 6.26×10^12^, error % = 3.03×10^-17^; friendliness: BF_10_ = 2.65×10^13^, error % = 2.28×10^-16^; intentionality: BF_10_ = 2.15×10^7^, error % = 9.62×10^-12^; abduction: BF_10_ = 7.23×10^6^, error % = 1.47×10^-11^). These results provide clear evidence of dimension-specific tuning: that the predictor corresponding to each dimension contributes uniquely to explaining ratings along that dimension.

**Figure S12.**
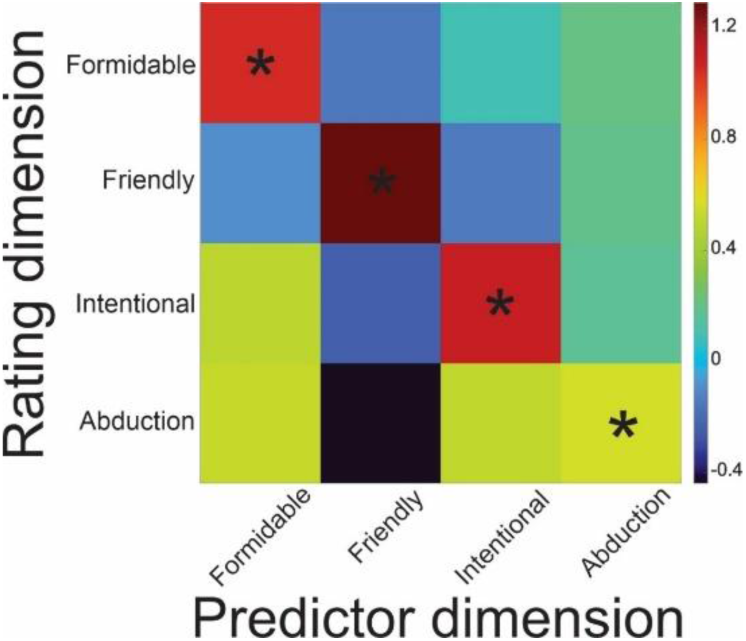
Group level beta matrix for the 4D model Note: Each cell denotes the mean regression coefficient relating a given rating dimension to a given PredF predictor. Rows – rating dimension (Dim1-Dim4); columns model predictors (PredF1–PredF4). The on-diagonal values are consistently largest in their respective rows, reflecting strong dimension-matching effects. Asterisks on the diagonal cells indicate dimensions where there was extreme evidence that the diagonal coefficients were larger than the mean off-diagonal coefficients.

## Experiment 3 – computational dissociation analysis for the 4D model

**Figure S13.**
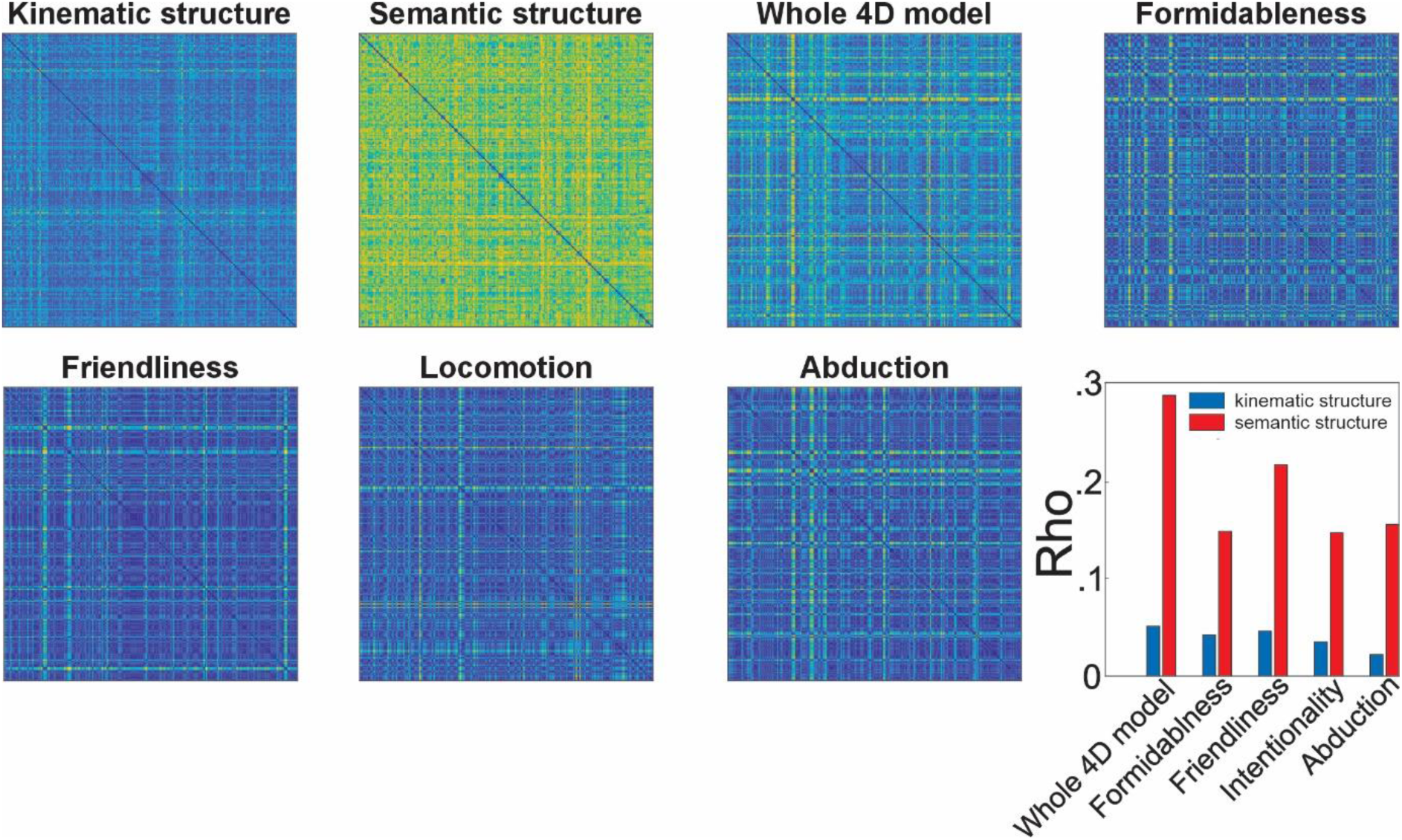
Representational dissimilarity matrices and Spearman rank correlations between kinematic and semantic models and the 4D action space model and individual dimensions Note: RDMs for kinematic and semantic structure models, the 4D action space model and its constituent dimensions. Correlations were all highly significant (all ps < .00001), and like the 5D model were much better correlated with the semantic (ρ = .287; ρ^2^ = 8.24%) than kinematic model (ρ = .051; ρ^2^ = .26%). Correlations of overlapping dimensions showed a similar profile of magnitudes across both models. Multiple regression RSA showed that the combined model (Kinematics + 4D Model) explained 11% of the total variance in action semantics, and the 4D model was a better predictor (β = .286) of action meaning than kinematics (β = .165). Partial correlation analysis showed that when controlling for kinematics, the relationship between 4D model and semantics remained mainly intact (Partial r_s_ = .287, p < .001 compared to ρ = .370).

## Notes

https://osf.io/cek59/overview

### Competing Interest Statement

The authors have declared no competing interest.

## References

Alaerts, K., Senot, P., Swinnen, S. P., Craighero, L., Wenderoth, N., & Fadiga, L. (2010). Force requirements of observed object lifting are encoded by the observer’s motor system: a TMS study. European Journal of Neuroscience, 31(6), 1144–1153. 10.1111/j.1460-9568.2010.07124.x

Altschul, D. M. (2024). Whither dominance? An enduring evolutionary legacy of primate sociality. Personality Neuroscience, 7, e1. 10.1017/pen.2023.13

Ansuini, C., Cavallo, A., Bertone, C., & Becchio, C. (2015). Intentions in the brain: the unveiling of Mister Hyde. Neuroscientist, 21(2), 126–135. 10.1177/1073858414533827

Anwyl-Irvine, A. L., Dalmaijer, E. S., Hodges, N., & Evershed, J. K. (2021). Realistic precision and accuracy of online experiment platforms, web browsers, and devices. Behavior research methods, 53, 1407–1425. 10.3758/s13428-020-01501-5

Anwyl-Irvine, A. L., Massonnié, J., Flitton, A., Kirkham, N., & Evershed, J. K. (2020). Gorilla in our midst: An online behavioral experiment builder. Behavior research methods, 52, 388–407. 10.3758/s13428-019-01237-x

Bailey, H. (2017). the OBS Project Contributors. In Open Broadcasting Software.

Barraclough, N. E., & Wightman, G. (2026). Testing a model of the cognitive structure of perceivable human actions. Visual Cognition, 1–19. 10.1080/13506285.2026.2636690

Becchio, C., Cavallo, A., Begliomini, C., Sartori, L., Feltrin, G., & Castiello, U. (2012). Social grasping: from mirroring to mentalizing. Neuroimage, 61(1), 240–248. 10.1016/j.neuroimage.2012.03.013

Becchio, C., Sartori, L., & Castiello, U. (2010). Towards you: the social side of actions. Current Directions in Psychological Science, 19(3), 183–188. 10.1177/0963721410370131

Bockes, A., Hebart, M. N., & Lingnau, A. (2025). Revealing Key Dimensions Underlying the Recognition of Dynamic Human Actions. Communications Psychology, 3(1), 149. 10.1038/s44271-025-00338-y

Carney, D. R., Hall, J. A., & LeBeau, L. S. (2005). Beliefs about the nonverbal expression of social power. journal of Nonverbal Behavior, 29(2), 105–123. 10.1007/s10919-005-2743-z

Chen, J., Paciocco, J. U., Deng, Z., & Culham, J. C. (2023). Human neuroimaging reveals differences in activation and connectivity between real and pantomimed tool use. Journal of Neuroscience, 43(46), 7853–7867.

Cole, E. J., Slocombe, K. E., & Barraclough, N. E. (2017). Abilities to explicitly and implicitly infer intentions from actions in adults with autism spectrum disorder. Journal of Autism and Developmental Disorders. 10.1007/s10803-017-3425-5

Darwin, C. (2025). The expression of the emotions in man and animals. In Death, Loss, Memory and Mourning in the Long Nineteenth Century, 1780–1914 (pp. 163–177). Routledge.

de Gelder, B. (2006). Towards the neurobiology of emotional body language. Nature Reviews Neuroscience, 7, 242–249. 10.1038/nrn1872

de Gelder, B., Van den Stock, J., Meeren, H. K. M., Sinke, C. B. A., Kret, M. E., & Tamietto, M. (2010). Standing up for the body. Recent progress in uncovering the networks involved in the perception of bodies and bodily expressions. Neurosci Biobehav Rev. 10.1016/j.neubiorev.2009.10.008

de la Rosa, S., Ferstl, Y., & Bulthoff, H. H. (2016). Visual adaptation dominates bimodal visual-motor action adaptation. Sci Rep, 6, 23829. 10.1038/srep23829

de la Rosa, S., Streuber, S., Giese, M., Bulthoff, H. H., & Curio, C. (2014). Putting actions in context: visual action adaptation aftereffects are modulated by social contexts. PLoS One, 9(1), e86502. 10.1371/journal.pone.0086502

de Lange, F. P., Spronk, M., Willems, R. M., Toni, I., & Bekkering, H. (2008). Complementary systems for understanding action intentions. Curr Biol, 18(6), 454–457. 10.1016/j.cub.2008.02.057

Decety, J., & Grezes, J. (1999). Neural mechanisms subserving the perception of human actions. Trends in Cognitive Sciences, 3(5), 172–178. 10.1016/S1364-6613(99)01312-1

Devlin, J., Chang, M.-W., Lee, K., & Toutanova, K. (2019). Bert: Pre-training of deep bidirectional transformers for language understanding. Proceedings of the 2019 conference of the North American chapter of the association for computational linguistics: human language technologies, volume 1 (long and short papers),

Dima, D. C., Tomita, T. M., Honey, C. J., & Isik, L. (2022). Social-affective features drive human representations of observed actions. Elife, 11, e75027. 10.7554/eLife.75027

Edelman, S. (1998). Representation is representation of similarities. Behavioral and brain sciences, 21(4), 449–467. 10.1017/S0140525X98001253

Edelman, S., & Intrator, N. (2003). Towards structural systematicity in distributed, statically bound visual representations. Cognitive Science, 27(1), 73–109. 10.1207/s15516709cog2701_3

Fedorov, L. A., Chang, D.-S., Giese, M. A., Bülthoff, H. H., & De la Rosa, S. (2018). Adaptation aftereffects reveal representations for encoding of contingent social actions. Proceedings of the National Academy of Sciences, 115(29), 7515–7520. 10.1073/pnas.1801364115

Ferstl, Y., Bülthoff, H., & de la Rosa, S. (2017). Action recognition is sensitive to the identity of the actor. Cognition, 166, 201–206. 10.1016/j.cognition.2017.05.036

Friston, K., Mattout, J., & Kilner, J. (2011). Action understanding and active inference. Biol Cybern, 104(1-2), 137–160. 10.1007/s00422-011-0424-z

Garcia, K., McMahon, E., Conwell, C., Bonner, M., & Isik, L. (2025). Modeling dynamic social vision highlights gaps between deep learning and humans. International Conference on Learning Representations,

Garcia, K., McMahon, E., Conwell, C., Bonner, M. F., & Isik, L. (2024). Dynamic, social vision highlights gaps between deep learning and human behavior and neural responses. Cognitive Computational Neuroscience,

Gärdenfors, P. (2004a). Conceptual spaces as a framework for knowledge representation. Mind and matter, 2(2), 9–27. 10.1017/S0140525X04280098

Gärdenfors, P. (2004b). Conceptual spaces: The geometry of thought. MIT Press.

Giese, M. A., & Poggio, T. (2003). Neural mechanisms for the recognition of biological movements. Nature Reviews Neuroscience, 4, 179–192. 10.1038/nrn1057

Indow, T., & Kanazawa, K. (1960). Multidimensional mapping of Munsell colors varying in hue, chroma, and value. Journal of experimental psychology, 59(5), 330. 10.1037/h0044796

JASP-Team. (2021). JASP (Version 0.16) [computer software]. In https://jasp-stats.org/

Jiang, M., Dong, J., Ma, D., Sun, J., He, J., & Lang, L. (2022). Inception spatial temporal graph convolutional networks for skeleton-based action recognition. 2022 International Symposium on Control Engineering and Robotics (ISCER),

Johansson, G. (1973). Visual perception of biological motion and a model for its analysis. Perception and Psychophysics, 14, 201–211. 10.3758/BF03212378

Jozwik, K. M., O’Keeffe, J., Storrs, K. R., Guo, W., Golan, T., & Kriegeskorte, N. (2022). Face dissimilarity judgments are predicted by representational distance in morphable and image-computable models. Proceedings of the National Academy of Sciences, 119(27), e2115047119. 10.1073/pnas.2115047119

Kabulska, Z., & Lingnau, A. (2023). The cognitive structure underlying the organization of observed actions. Behavior research methods, 55(4), 1890–1906. 10.3758/s13428-022-01894-5

Koo, T. K., & Li, M. Y. (2016). A guideline of selecting and reporting intraclass correlation coefficients for reliability research. Journal of chiropractic medicine, 15(2), 155–163. 10.1016/j.jcm.2016.02.012

Koppensteiner, M. (2013). Motion cues that make an impression: Predicting perceived personality by minimal motion information. J Exp Soc Psychol, 49(6), 1137–1143. 10.1016/j.jesp.2013.08.002

Koppensteiner, M., Stephan, P., & Jäschke, J. P. M. (2016). Moving speeches: Dominance, trustworthiness and competence in body motion. Personality and Individual Differences, 94, 101–106. 10.1016/j.paid.2016.01.013

Kriegeskorte, N., & Kievit, R. A. (2013). Representational geometry: integrating cognition, computation, and the brain. Trends in Cognitive Sciences, 17(8), 401–412. 10.1016/j.tics.2013.06.007

Kriegeskorte, N., Mur, M., & Bandettini, P. A. (2008). Representational similarity analysis-connecting the branches of systems neuroscience. Front Syst Neurosci, 2, 249. 10.3389/neuro.06.004.2008

Landauer, T. K., & Dumais, S. T. (1997). A solution to Plato’s problem: The latent semantic analysis theory of acquisition, induction, and representation of knowledge. Psychological review, 104(2), 211–240. 10.1037/0033-295X.104.2.211

McMahon, E., Bonner, M. F., & Isik, L. (2023). Hierarchical organization of social action features along the lateral visual pathway. Current biology, 33(23), 5035–5047. e5038. 10.1016/j.cub.2023.10.015

Moore, J., & Haggard, P. (2008). Awareness of action: Inference and prediction. Consciousness and cognition, 17(1), 136–144. 10.1016/j.concog.2006.12.004

Nosofsky, R. M. (1986). Attention, similarity, and the identification–categorization relationship. Journal of Experimental Psychology: General, 115(1), 39. 10.1037/0096-3445.115.1.39

Nosofsky, R. M. (1991). Tests of an exemplar model for relating perceptual classification and recognition memory. Journal of experimental psychology: human perception and performance, 17(1), 3. 10.1037//0096-1523.17.1.3

Runeson, S., & Frykholm, G. (1981). Visual perception of lifted weight. Journal of experimental psychology: human perception and performance, 7, 733–740. 10.1037/0096-1523.7.4.733

Sartori, L., Becchio, C., & Castiello, U. (2011). Cues to intention: the role of movement information. Cognition, 119(2), 242–252. 10.1016/j.cognition.2011.01.014

Saxe, R., & Kanwisher, N. (2003). People thinking about thinking people The role of the temporo-parietal junction in “theory of mind”. Neuroimage, bras19(4), 1835–1842. 10.1016/S1053-8119(03)00230-1

Shepard, R. N. (1962a). The analysis of proximities: multidimensional scaling with an unknown distance function. I. Psychometrika, 27(2), 125–140. 10.1007/BF02289630

Shepard, R. N. (1962b). The analysis of proximities: Multidimensional scaling with an unknown distance function. II. Psychometrika, 27(3), 219–246. 10.1007/BF02289621

Shepard, R. N. (1982). Geometrical approximations to the structure of musical pitch. Psychological review, 89(4), 305. 10.1037/0033-295X.89.4.305

Shepard, R. N. (1987). Toward a universal law of generalization for psychological science. Science, 237(4820), 1317–1323. 10.1126/science.3629243

Shepard, R. N., Hovland, C. I., & Jenkins, H. M. (1961). Learning and memorization of classifications. Psychological monographs: General and applied, 75(13), 1. 10.1037/h0093825

Spunt, R. P., Satpute, A. B., & Lieberman, M. D. (2011). Identifying the what, why, and how of an observed action: an fMRI study of mentalizing and mechanizing during action observation. Journal of cognitive neuroscience, 23(1), 63–74.

Stolier, R. M., Hehman, E., Keller, M. D., Walker, M., & Freeman, J. B. (2018). The conceptual structure of face impressions. Proceedings of the National Academy of Sciences, 115(37), 9210–9215. 10.1073/pnas.1807222115

Tarhan, L., & Konkle, T. (2020). Sociality and interaction envelope organize visual action representations. Nature communications, 11(1), 3002. 10.1038/s41467-020-16846-w

Thoresen, J. C., Vuong, Q. C., & Atkinson, A. P. (2012). First impressions: gait cues drive reliable trait judgements. Cognition, 124(3), 261–271. 10.1016/j.cognition.2012.05.018

Thornton, M. A., & Tamir, D. I. (2020). People represent mental states in terms of rationality, social impact, and valence: Validating the 3d Mind Model. Cortex, 125, 44–59. 10.1016/j.cortex.2019.12.012

Troje, N. F. (2002). Decomposing biological motion: A framework for analysis and synthesis of human gait patterns. Journal of Vision, 2(5), 371–387. 10.1167/2.5.2

Troje, N. F., Sadr, J., Geyer, H., & Nakayama, K. (2006). Adaptation aftereffects in the perception of gender from biological motion. Journal of Vision, 6, 850–857. 10.1167/6.8.7

Troje, N. F., Westhoff, C., & Lavrov, M. (2005). Person identification from biological motion: Effects of structural and kinematic cues. Perception & Psychophysics, 67(4), 667–675. 10.3758/BF03193523

Tucciarelli, R., Wurm, M., Baccolo, E., & Lingnau, A. (2019). The representational space of observed actions. Elife, 8, e47686. 10.7554/eLife.47686

Valentine, T. (1991). A unified account of the effects of distinctiveness, inversion, and race in face recognition. The Quarterly Journal of Experimental Psychology Section A, 43(2), 161–204. 10.1080/14640749108400966

Van Overwalle, F., & Baetens, K. (2009). Understanding others’ actions and goals by mirror and mentalizing systems: a meta-analysis. Neuroimage, 48(3), 564–584. 10.1016/j.neuroimage.2009.06.009

Vinton, L. C., Preston, C., de la Rosa, S., Mackie, G., Tipper, S. P., & Barraclough, N. E. (2023). Four fundamental dimensions underlie the perception of human actions. Attention, Perception, & Psychophysics, 1–23. 10.3758/s13414-023-02709-1

Vlasceanu, A. M., de la Rosa, S., & Barraclough, N. E. (2024). Perceptual discrimination of action formidableness and friendliness and the impact of autistic traits. Scientific reports, 14(1), 25554. 10.1038/s41598-024-76488-6

Wiggins, J. S. (1979). A psychological taxonomy of trait-descriptive terms: The interpersonal domain. Journal of Personality and Social Psychology, 37(3), 395. 10.1037/0022-3514.37.3.395

Wurm, M. F., Ariani, G., Greenlee, M. W., & Lingnau, A. (2015). Decoding Concrete and Abstract Action Representations During Explicit and Implicit Conceptual Processing. Cereb Cortex. 10.1093/cercor/bhv169

Wurm, M. F., & Caramazza, A. (2019). Lateral occipitotemporal cortex encodes perceptual components of social actions rather than abstract representations of sociality. Neuroimage, 202, 116153. 10.1016/j.neuroimage.2019.116153

Wurm, M. F., & Caramazza, A. (2021). Two ‘what’pathways for action and object recognition. Trends in Cognitive Sciences.

Wurm, M. F., Caramazza, A., & Lingnau, A. (2017). Action categories in lateral occipitotemporal cortex are organized along sociality and transitivity. Journal of Neuroscience, 37(3), 562–575. 10.1523/JNEUROSCI.1717-16.2016

Yan, S., Xiong, Y., & Lin, D. (2018). Spatial temporal graph convolutional networks for skeleton-based action recognition. Proceedings of the AAAI conference on artificial intelligence,

